# Comparative Evaluation of Deep Generative Models for Capturing Topological Features in Brain Structural Connectivity Matrices

**DOI:** 10.64898/2026.06.03.729714

**Authors:** Chisato Kumada, Tomoyuki Hiroyasu, Satoru Hiwa

## Abstract

Structural connectivity (SC) data are crucial for brain network analysis, but SC-based machine learning often suffers from limited data availability, hindering model generalization and robustness. Although data augmentation using deep generative models has attracted increasing attention, it remains unclear how different models capture the complex topological features of SC data. To clarify the model-specific generative characteristics of deep generative models for ROI-level weighted SC matrix generation, this study compares three representative models: variational autoencoder (VAE), Wasserstein generative adversarial network with gradient penalty (WGAN-GP), and denoising diffusion probabilistic models (DDPM). We systematically evaluated these models using both synthetic datasets as diagnostic benchmarks to isolate model-specific biases in reproducing graph properties and two real-world SC datasets. Generation quality was assessed using graph-theoretic metric comparisons and visual inspection of the generated adjacency matrices across four training runs. For the empirical SC datasets, we also evaluated inter-subject variability and sample diversity. WGAN-GP performed consistently across synthetic datasets and achieved the strongest results on most metrics for both SC datasets, most closely reproducing variability and diversity. VAE captured average characteristics but underestimated sample variability. DDPM reproduced stochastic connection patterns but struggled with coordinate-specific patterns and run-to-run consistency. Despite WGAN-GP’s balanced performance, each model had distinct strengths and limitations, emphasizing the need to consider relevant graph properties when comparing generative models. All models struggled to reproduce strict global constraints, such as planarity. These findings highlight the importance of evaluating structural constraints alongside statistical similarity and accounting for graph-specific properties when developing generative models.

## 1. Introduction

Recently, brain structural connectivity (SC) has gained significant attention in neuroscience research due to its potential to reveal insights into brain function and organization.^3,4^ SC can be represented as an adjacency matrix, in which rows and columns correspond to brain regions and each edge represents the connectivity between a pair of regions. There has been a growing interest in applying SC to various applications, including investigations into brain development and aging, and analysis of neurological and brain disorders.^5^ Specifically, machine learning approaches based on SC have been utilized to diagnose and understand disorders such as Alzheimer’s disease^6^ and multiple sclerosis.^7^ However, the practical application of SC-based models often suffers from limited data availability, which affects the generalization and robustness of machine learning approaches. Acquiring large-scale diffusion MRI (dMRI) datasets is often challenging due to high costs and time-consuming data collection processes.^8^ Neuroimaging studies frequently suffer from limited sample sizes, which introduces bias in training data and limits the generalizability of machine learning models.^9^ Such data scarcity often leads to overfitting and unreliable model performance in the downstream machine learning analyses, such as prediction or diagnosis.

Although data augmentation is an effective strategy for addressing data scarcity, traditional augmentation methods such as rotation or flipping are not applicable to SC matrices. Because SC matrices are represented as an adjacency matrix where each row and column corresponds to a specific brain region, such geometric transformations would destroy this correspondence. Therefore, generating plausible SC data requires approaches that can preserve its topological characteristics. Brain structural networks are characterized by complex topological features, such as short path length, high clustering coefficient, hub presence, and modularity, alongside functionally critical, strong edge weights.^3,10^ These features fundamentally facilitate the efficient segregation and integration of information. Thus, generating biologically plausible SC data requires capturing the high-dimensional topological dependencies among multiple regions, a task that is inherently challenging for simple approaches.

To overcome this limitation, deep generative models have been explored for SC data augmentation, though a comprehensive understanding of their comparative capabilities remains lacking. By learning the underlying distribution of the training data, these models generate new samples that resemble the original data. Representative models such as variational autoencoders (VAE)^11^ and generative adversarial networks (GANs)^12^ have been successfully applied to generate synthetic SC data, demonstrating clear improvements in downstream tasks.^13–15^ Additionally, diffusion models^16^ have emerged as a promising approach for generating high-quality synthetic data across various domains. While their direct application to SC data remains underexplored, researchers have begun exploring diffusion models for brain connectivity, such as functional connectivity (FC) generation and bidirectional SC-FC mapping,^17,18^ making diffusion models an important emerging candidate for SC generation as well. However, previous studies have primarily focused on individual generative models for SC data augmentation, using different datasets and evaluation metrics. This makes it challenging to directly compare the ability of different generative models to capture the graph topology in this context. Given that VAEs, GANs, and diffusion models rely on fundamentally different generative paradigms, each model is expected to exhibit distinct behaviors when reconstructing SC structures. Addressing this gap and revealing how each model learns brain network characteristics is a crucial step toward understanding the fundamental capabilities and limitations of deep generative models in this domain.

To elucidate how deep generative models learn and reproduce the topological features of SC data, this study conducts a systematic empirical comparison of three representative implementations of deep generative models: VAE, Wasserstein GAN with gradient penalty (WGAN-GP),^19^ and DDPM. In these comparative evaluations, we focused not only on overall generation quality but also on which graph properties each model reproduced well or poorly. For this purpose, we firstly used five synthetic graph datasets with controlled characteristics, representing hub structure, modularity, small-worldness, weighted edges, and strict global constraints. Then we used two empirical SC datasets, the NIMH Healthy Volunteer Dataset and Human Connectome Project Young Adult, in which these properties coexist. In this study, SC generation refers to generating ROI-level weighted connectivity matrices defined on a fixed anatomical parcellation. Model performance was evaluated using Maximum Mean Discrepancy (MMD)-based distributional comparisons, dataset-specific graph-theoretical metrics, and visual inspection of generated adjacency matrices across four runs. In addition, we evaluated inter-subject variation and sample diversity for SC datasets. By evaluating these models under a common framework across controlled synthetic graphs and empirical SC datasets, this study systematically identifies their capabilities and limitations in reproducing different graph properties. These findings provide an empirical basis for selecting and improving generative models for SC generation tasks.

## 2. Methods

In this study, we investigated the performance of three generative models for augmenting brain SC data. To comprehensively evaluate performance against SC data, this study employed both synthetic and real-world SC datasets. Since real-world SC data is complex and lacks explicitly defined target structures, evaluating solely on real SC data makes it difficult to identify which structural features the model struggles to reproduce. To address this, our evaluation framework first assesses the ability to reproduce individual graph properties with known network characteristics. Subsequently, we evaluate the models on real-world SC data, where multiple structural properties coexist.

An overview of the experimental framework used in this study is illustrated in Figure 1, which includes the datasets, the compared models, and the evaluation protocols.

**Figure 1.**
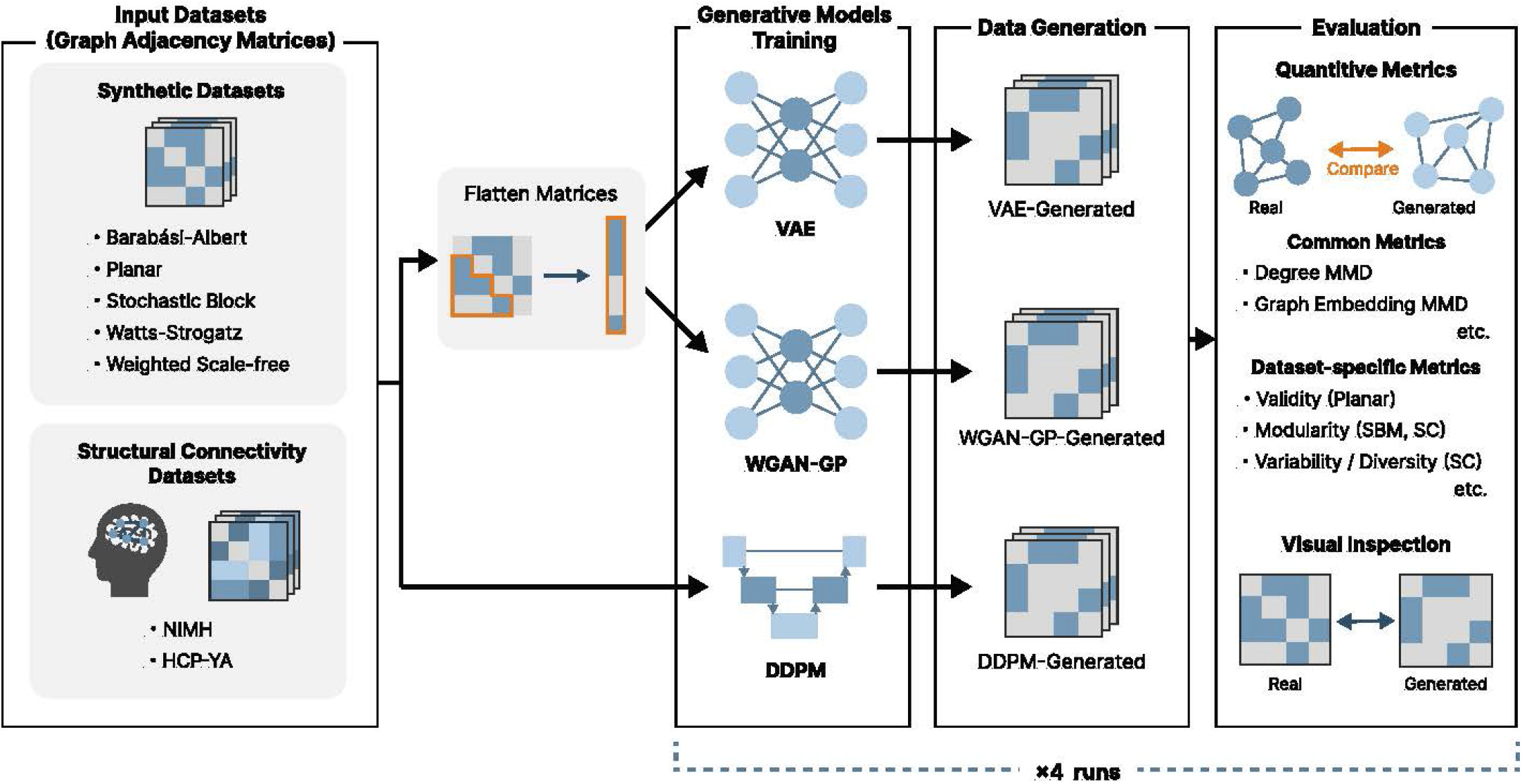
Overview of the overall methodology used in this study.

### 2.1. Compared Models

To evaluate the potential of different generative models, we compared the performance of three fundamental generative models: VAE, WGAN-GP, and DDPM. Details of each model are described below. Detailed loss functions and model architectures can be found in the Supplementary Information.

#### 2.1.1. Variational Autoencoder

VAE, proposed by Kingma and Welling,^11^ is a generative model that learns to encode input data into a latent space and decode samples from this space back to the data space. The model consists of two neural networks: an encoder and a decoder. The encoder maps the input data *x* into a latent representation *z*, while the decoder reconstructs the original data from the latent representation. The model is trained to minimize the sum of a reconstruction loss and a Kullback-Leibler (KL) divergence loss.

In this study, both the encoder and the decoder were fully connected neural networks that took the flattened lower-triangular part of the adjacency matrix as input and produced the output.

#### 2.1.2. Wasserstein GAN with Gradient Penalty

WGAN-GP is a variant of GANs that improves training stability by using the Wasserstein distance as a loss function and adding a gradient penalty term, as proposed by Arjovsky et al.^20^ and further refined by Gulrajani et al.^19^ The model consists of two neural networks: a generator and a critic. The generator creates synthetic data from random noise, while the critic evaluates the authenticity of the given data, which can be either real or generated.

In this study, both the generator and the critic were fully connected neural networks that processed the flattened lower-triangular part of the adjacency matrix.

#### 2.1.3. Denoising Diffusion Probabilistic Models (DDPM)

DDPM, proposed by Ho et al.,^16^ is a generative model that learns to convert random noise into data through a series of denoising steps. The model has two processes: a forward diffusion process and a reverse denoising process. In the forward process, Gaussian noise is added to the data over a series of time steps. A neural network is trained to predict the noise added to the original data, given the noisy data and the time step. In the reverse process, the model generates data from random noise by iteratively removing the predicted noise.

For the implementation of the DDPM, we adopted the standard U-Net^21^ architecture, which is commonly used in diffusion models. Because VAE and WGAN-GP used MLP-based architectures, the effects of network architecture and generative paradigm cannot be fully separated. The number of the noising and denoising steps was set to *T* = 1000 as in the original paper.

### 2.2. Datasets

Generating realistic SC data requires reproducing topological features inherent in brain networks, such as short path lengths, high clustering coefficients, hub presence, and modularity.^3^ These features are crucial for brain function, since they fundamentally facilitate the efficient segregation and integration of information. Specifically, small-worldness, which refers to the coexistence of short path lengths and high clustering coefficients, balances specialized local processing with effective global communication at a lower cost.^22^ Modularity, which refers to the organization of the network into densely connected clusters with sparse connections between them, supports specialized regional processing. Hubs act as central nodes for information exchange, guaranteeing an optimal balance between this segregation and integration. Furthermore, preserving strong edge weights is also important, as they serve as the foundational backbone that constrains and supports complex communication processes within the brain.^10^

Reproducing these features is therefore essential for generating biologically plausible SC data. To systematically evaluate the models’ ability to reproduce different graph properties under controlled conditions, we selected five distinct synthetic datasets as diagnostic benchmarks.

#### 2.2.1. Synthetic Datasets

We selected five types of synthetic datasets to reproduce a range of graph structures and properties. The size of the synthetic datasets was fixed at 64 nodes. Detailed generation procedures and parameter settings for each synthetic dataset are provided in the Supplementary Information.

##### Barabási-Albert (*N* = 500)

We utilized the BA dataset to investigate the reproduction of specific degree distributions and hub nodes. BA models generate scale-free networks featuring few high-degree hubs and many low-degree nodes. Although brain networks exhibit an exponential degree distribution rather than strictly scale-free degree distribution,^23,24^ we selected the BA dataset because its ability to reproduce hub structures and degree distributions is highly relevant to capturing the fundamental properties of SC.

##### Planar (*N* = 200)

The planar dataset serves as a benchmark for evaluating the ability to learn global topological constraints. Planar graphs have a strict global rule that edges must not cross in 2-dimensional space; therefore, investigating whether the generated graphs meet planarity constraints gives insights into a model’s capacity to learn implicit global rules. We used an existing benchmark dataset provided by Martinkus et al^1^, which contains fewer samples (*N* = 200) than the other synthetic datasets (*N* = 500).

##### Stochastic Block Model (*N* = 500)

The SBM dataset was selected to assess the model’s ability to reproduce community structure. SBM graphs consist of distinct communities with random intra- and inter-connections, making them suitable for evaluating community reproduction. Since the brain has an explicit modularity, evaluating the reproduction of these structures is critical for SC generation.

##### Watts-Strogatz (*N* = 500)

The WS dataset was chosen to evaluate the models’ capability to capture small-worldness. Because small-worldness is a fundamental property of brain networks that enables efficient information transfer, assessing its reproducibility is essential for validating generative models.

##### Weighted Scale-Free (*N* = 500)

We selected a weighted scale-free network to assess the models’ capability to generate weighted graphs. Since SC data forms a weighted network, generating not only the correct binary structure but also precise edge weights is important for reflecting the foundational backbone. We generated 500 weighted scale-free graphs using the method proposed by Yook et al.^25^

#### 2.2.2. Structural Connectivity Datasets

To evaluate the generative models on complex real-world brain connectivity data, we utilized two empirical SC datasets: the NIMH Healthy Volunteer Dataset and the Human Connectome Project Young Adult (HCP-YA) S1200 Dataset. SC matrices were reconstructed from diffusion MRI data using closely matched tractography procedures. Tractography was performed using the iFOD2 algorithm,^26^ followed by SIFT2.^27^ The whole brain was parcellated into 116 regions based on the automated anatomical labeling (AAL) atlas.^28^ SC matrices were constructed by summing the SIFT2 weights of streamlines connecting each pair of regions, scaled by the inverse node volumes. Detailed reconstruction procedures are provided in the Supplementary Information.

##### The NIMH Healthy Volunteer Dataset (N=107)

The dataset was derived from diffusion MRI and T1-weighted MRI data of healthy adult participants, originally obtained from the NIMH Healthy Volunteer Dataset (https://doi.org/10.18112/openneuro.ds004215.v1.0.0).^2^ In total, we used data from 107 participants for our experiments.

##### HCP-YA S1200 Dataset (N=1,065)

We additionally used the Human Connectome Project Young Adult (HCP-YA) S1200 release to evaluate the models on a larger SC dataset. Participants without diffusion-weighted imaging data were excluded from the analysis, resulting in a final sample of 1,065 participants.

### 2.3. Training Setup

We evaluated the performance of the three generative models on the seven datasets described above. Each model was trained separately on each of the five synthetic datasets and on two SC datasets.

#### 2.3.1. Data Preprocessing

To reflect the fixed node correspondence of SC matrices, we applied deterministic node ordering to the synthetic datasets. In SC matrices, each node corresponds to a specific brain region, and this correspondence is consistent across samples. Accordingly, our synthetic datasets were designed to evaluate the reproduction of controlled topological properties in fixed-coordinate adjacency matrix generation. Specifically, we applied the following methods: depth-first search (DFS) for the BA, Planar, and WSF datasets; k-core ordering for the WS dataset; and community-based k-core ordering for the SBM dataset.

Regarding edge features, SC edge weights were calculated from SIFT2-weighted streamlines with inverse node-volume scaling, as described in Section 2.2.2. We treated these volume-corrected weights as the target representation for generation and did not apply additional normalization before model training. While we acknowledge that generative models typically benefit from input data being scaled to a specific optimal range for stable training, we prioritized preserving the inherent distribution and variance of the volume-corrected connectivity data.

#### 2.3.2. Data Splitting and Assessment of Run-to-Run Variability

To assess the run-to-run variability in generative performance arising from different training/validation splits and random initializations, we conducted four training runs for each model-dataset combination.

First, we split each dataset into a discovery set and a test set. The test set comprised 20% of the samples for the synthetic datasets and the HCP dataset, and 25% for the NIMH dataset. The test set was fixed across all runs for final model evaluation and held out from model training and hyperparameter optimization.

Second, the discovery set was divided into four folds. Across the four runs, each fold was used as the validation set exactly once, while the remaining three folds were used as the training set. The validation fold was rotated across the four runs so that each fold was used for validation exactly once. A different random seed was also assigned to each run, and the same set of four seeds was used across all model-dataset combinations.

A detailed dataset separation procedure is provided in the Supplementary Information.

#### 2.3.3. Model Training Details

All models were implemented in PyTorch^31^ and trained with the Adam optimizer. Hyperparameters were optimized using Optuna^32^ in the first run for each model-dataset combination. The hyperparameter values selected in the first run were fixed and reused for runs 2-4. For both hyperparameter optimization and early stopping, validation loss was used for VAE and DDPM, whereas validation MMD^33^ was used for WGAN-GP because adversarial loss does not necessarily reflect the quality of generated samples. Detailed descriptions of the computational environment, training settings, and hyperparameter search spaces are provided in the Supplementary Table S1. The selected hyperparameters for each model and dataset are summarized in Supplementary Tables S2 and S3.

After training, we generated 1000 samples from each model using its standard generative procedure. For DDPM, sampling was performed in batches of 500 samples for synthetic datasets and 250 samples for SC datasets due to limited GPU memory.

#### 2.3.4. Computational Cost

Training time, peak GPU memory usage, and sampling time were measured for runs 2-4. Training and sampling times were measured as wall-clock time. Run 1 was excluded because comparable measurements were not available. The number of trainable parameters and the number of training epochs were also recorded across all runs.

### 2.4. Evaluation Metrics

To evaluate how generative models learn the aforementioned topological features essential for brain networks, we selected the following evaluation metrics. The evaluation included both metrics common to all datasets and metrics defined specifically for individual datasets.

#### 2.4.1. Edge Thresholding

For the synthetic datasets, all graphs were thresholded to achieve an edge density similar to that of the training dataset before evaluation, since some models tended to generate subtle edges rather than no edges. Thresholding was therefore applied to compare topological fidelity under similar sparsity conditions. The threshold value was searched in increments of 0.01 to minimize the difference in edge density between the generated graphs and the corresponding training dataset. Non-weighted graphs were binarized after thresholding. The selected threshold for each run, the densities of the generated graphs before and after thresholding, and the density of the corresponding training dataset are reported in the supplementary information.

For the SC datasets, we applied percentile-based thresholding to the edge weights of the generated graphs. This approach was employed to focus on the important connections in the brain network. We retained the top 5%, 15%, and 30% of the strongest edges in each generated graph for most evaluation metrics. However, for the strong edge preservation metric, we retained the top 1%, 5%, and 10% of the strongest edges to assess the models’ ability to reconstruct the most significant connections. For the hub preservation metric, no thresholding was applied, as node strength can be calculated without thresholding.

#### 2.4.2. Common Metrics

##### MMD-based Evaluation

Following previous studies,^1,34^ we used MMD to quantitatively assess the similarity between the test and generated datasets. Because MMD measures the distance between two underlying probability distributions based on finite samples, it allows us to evaluate the overall characteristics of statistical graph properties rather than relying on a single representative value. Specifically, we computed the MMD for the following graph-theoretical features: node degree, clustering coefficient, orbit counts, the spectrum of the graph Laplacian matrix, and graph embeddings.

Since MMD values are relative and depend on the specific datasets and features being compared, we calculated reference MMD values to provide context for interpreting the results. For the BA, SBM, WS, and WSF datasets, we generated new graphs using the same generation process as the original dataset to compute reference MMD values. For the planar and SC datasets, the reference values were calculated using the training and validation sets, since their generation processes were not explicitly defined.

##### Mean adjacency matrix visualization

Mean adjacency matrix visualization was performed to provide an overview of the generated graphs. The mean adjacency matrix was calculated by averaging the adjacency matrices of all graphs in each dataset generated by each model across all runs. This visual inspection was adopted because global quantitative metrics may sometimes overlook spatial biases or the disappearance of specific structural patterns.

#### 2.4.3. Dataset-specific Metrics

To evaluate whether the generative models successfully captured the intrinsic topological properties unique to each dataset, we employed specific metrics that quantify their essential structural characteristics.

##### Validity

For the Planar dataset, the validity of the generated graphs was evaluated by whether they had a planar structure, that is, the absence of edge crossings in 2-dimensional space. It was calculated to determine whether the generative models could implicitly learn the dataset’s global structural constraints.

##### Modularity

Modularity was calculated to assess community structure, a feature of both the SBM and SC datasets. Modularity quantifies the degree to which a network is partitioned into distinct clusters, with internal edge density higher than external edge density.

We detected communities using Clauset-Newman-Moore greedy modularity maximization, calculated the modularity of the detected communities, and compared the modularity distributions between the generated and test datasets by computing the Wasserstein distance between them.

##### Small-worldness

Since the WS and SC datasets are characterized by small-world properties, the small-worldness coefficient *σ* of each graph was calculated, following the definition proposed by Humphries and Gurney.^35^ This metric was chosen to determine if the generative models can reproduce the balance between local dense connections and efficient global paths, which are characteristics of brain networks. We compared the distributions of *σ* values between the generated and test datasets and calculated the Wasserstein distance between them.

##### Node Strength MMD

For the WSF and SC datasets, MMD of node strength was calculated to evaluate the models’ ability to reproduce the weighted graph. Node strength, defined as the sum of the weights of all edges connected to a node, is a pivotal metric that integrates topological connectivity with weight intensity. Evaluating node strength reveals the structural rules governing how weights are distributed across the network, such as the concentration of high-intensity connections on specific functional hubs.

##### Strong Edge/Hub Preservation

To further evaluate the structural fidelity of the generated SC matrices, we focused on how well strong edges and hubs prominent in the group-mean SC matrices are preserved. By focusing on the mean adjacency matrix, we assess whether the models can reproduce prominent connectivity patterns at the group level.

For strong edge reconstruction, we identified the top *n*% (*n* = 1, 5, 10) strongest edges in the mean adjacency matrix of the test dataset and compared them with those of the generated graphs using the Jaccard index. This indicates whether the model captures the stable, high-intensity structural backbones observed in the tractography-derived SC data. Regarding hub preservation, we calculated the Spearman correlation coefficient between the node strengths of the mean test dataset and those of the generated datasets. High correlation indicates that the model correctly identifies the hubs.

##### Edge Weight Variability

To evaluate whether the generated SC samples preserved inter-subject variability in edge weights, we calculated the standard deviation of each edge across samples for both the NIMH and HCP datasets. For each edge, the variance observed in the generated samples was compared with that in the corresponding test set.

##### Pairwise Distance and Nearest-Neighbor Analysis

For the NIMH and HCP SC datasets, we performed nearest-neighbor and pairwise distance analyses using the L2 distance between flattened lower-triangular elements of the adjacency matrices.

To evaluate the diversity of the generated samples, we calculated pairwise distances between all pairs of generated samples within each run, and performed the same analysis for the test set. The resulting pairwise distance distributions were compared between the generated samples from each model and the corresponding test set.

To further assess whether the generative models memorized the training data, we performed a nearest-neighbor analysis using the L2 distance between flattened adjacency matrices. For each generated sample, we calculated the distance to its nearest sample in the training set. The same analysis was performed for each test SC sample relative to the training set. The distributions of generated-to-training and test-to-training nearest-neighbor distances were then compared.

## 3. Results

### 3.1. Synthetic Datasets

We qualitatively evaluated the generated results for synthetic datasets using mean adjacency matrices (Figure 2), supplemented by individual adjacency matrices (Figure S1). All models exhibited residual differences from the target matrices, although their spatial distribution varied across models. The predominantly positive residuals in WSF indicated a tendency to underestimate edge weights across all three models.

**Figure 2.**
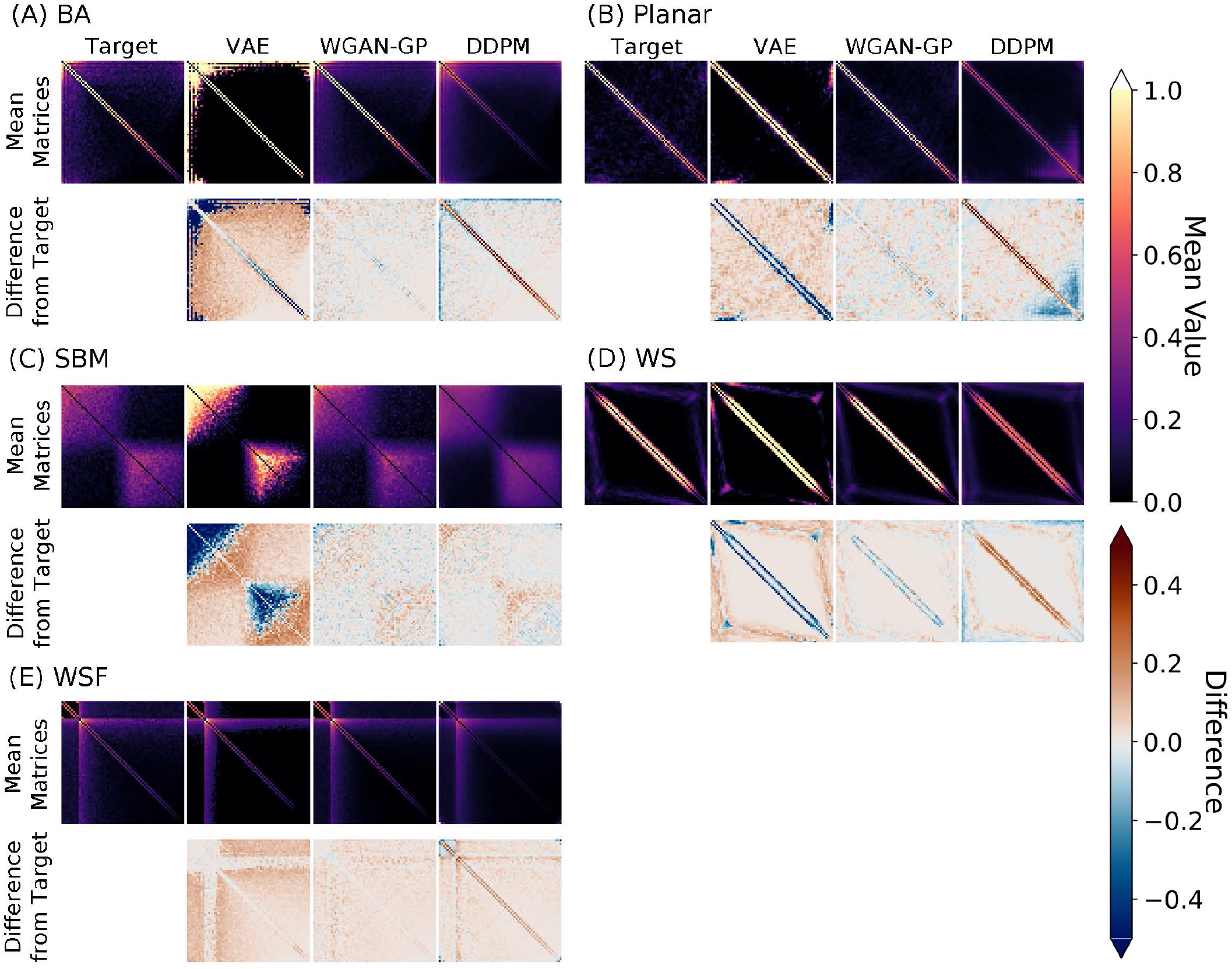
Mean adjacency matrix analysis across five synthetic datasets. (A-E). Top row: Mean adjacency matrices of the test set (target) and the generated graphs. For each model, the mean matrix was calculated by pooling all generated samples across runs. The color bar indicates the magnitude of connectivity. Lighter colors indicate greater values. Bottom row: Corresponding residual matrices (target - generated). Red indicates positive residuals, while blue indicates negative residuals. *Abbreviations*: VAE = variational autoencoder, WGAN-GP = Wasserstein GAN with gradient penalty, DDPM = denoising diffusion probabilistic models, BA = Barabási-Albert, SBM = stochastic block model, WS = Watts–Strogatz, WSF = weighted scalefree.

VAE replicated high-probability patterns (e.g., edges near the diagonal lines in BA and Planar) but exhibited a bias towards over-representing common patterns, including hub-related edges in BA and intra-community edges in SBM, as indicated by negative residuals. In contrast, low-probability edges, such as off-diagonal edges in BA and Planar, showed large positive residuals. VAE also tended to generate individual samples that resembled the mean adjacency matrices (Figure S1).

WGAN-GP showed a balanced ability to reproduce common patterns and generate random edges, capturing clear patterns without overemphasizing common edges, as observed in VAE. While WGAN-GP generated more visually plausible graphs with generally less pronounced residuals than the other models, continuous structures in the Planar and WS datasets were fragmented into scattered edges.

DDPM, in contrast, excelled at reproducing stochastic connection patterns, especially in SBM. While it also captured some of the fixed structures of the Planar, WS, and WSF, these structures were generally less pronounced than those in the test data, indicating incomplete reproduction of high-probability edges. Furthermore, comparisons across training runs (Figure S1) revealed substantial variation in preserving these structures. Notably, run 1, in which hyperparameter optimization was performed, successfully reproduced the fixed patterns.

Quantitative results showed that model performance depended strongly on the structural properties of the target graphs (Figure 3). All models struggled with datasets characterized by strict structural constraints. In the BA datasets, all models showed high MMD values for degree distributions, indicating difficulty in reproducing the scale-free property of BA. Validity scores for Planar graphs were universally low, and no model achieved the target small-worldness (*σ*) for WS graphs. Notably, across all datasets and metrics, no model consistently reached the performance level of the reference dataset. The results for each individual run, including MMD, planarity, and Wasserstein distance, are provided in Supplementary Tables S7 and S8.

**Figure 3.**
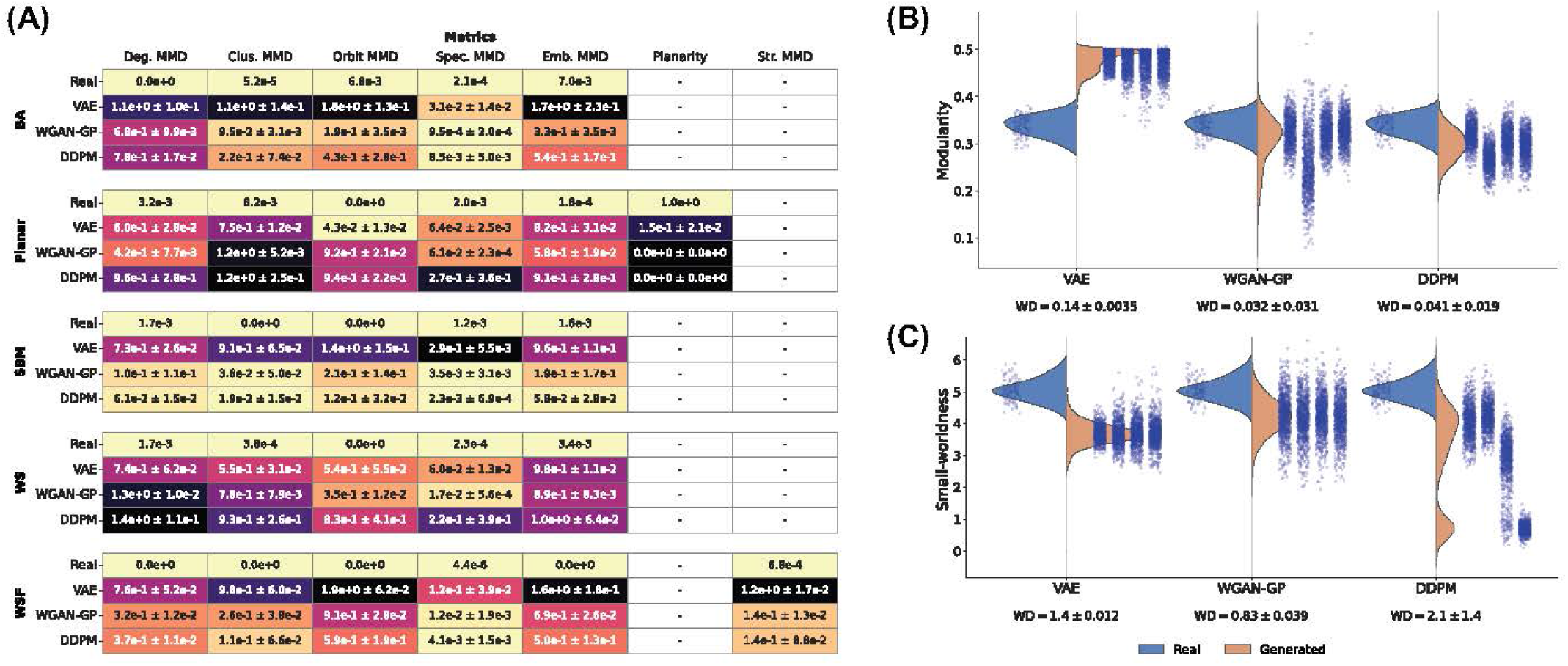
Quantitative evaluation results of generative models across synthetic datasets. (A) Heatmap of MMD and planarity results for synthetic datasets. Cells are colored based on the mean value across all models and datasets, with lighter colors indicating better performance. (B) Distributions of modularity scores and Wasserstein distances between generated graphs and the test set for the SBM dataset. (C) Distributions of *σ* values (small-worldness) and Wasserstein distances between generated graphs and the test set for the WS dataset. In (B) and (C), violin plots show distributions pooled across four independent runs, with scatter points representing individual runs, arranged from left to right in the order of runs 1-4. MMD and Wasserstein distance values are reported as mean ± standard deviation across four independent runs. *Abbreviations*: VAE = variational autoencoder, WGAN-GP = Wasserstein GAN with gradient penalty, DDPM = denoising diffusion probabilistic models, BA = Barabási-Albert, SBM = stochastic block model, WS = Watts–Strogatz, WSF = weighted scale-free.

VAE generally showed higher MMD values than the other models on the BA, SBM, and WSF datasets, although it was the only model that generated valid planar graphs in the planar dataset. WGAN-GP demonstrated relatively consistent performance across metrics, achieving lower MMD values than the other models for most metrics. It also achieved the lowest Wasserstein distances for modularity in SBM and small-worldness in WS. DDPM showed substantial variation in performance across datasets. It performed relatively well on the SBM and WSF datasets but struggled on the Planar and WS datasets. Notably, DDPM achieved the lowest MMDs in all MMD metrics for SBM.

### 3.2. Structural Connectivity Datasets

For both SC datasets, the mean matrices generated by VAE and WGAN-GP reproduced the major spatial patterns observed in the test data. DDPM-generated matrices were blurred in NIMH, whereas SC patterns were more visible in HCP (Figure 4). Individual adjacency matrices presented in Supplementary Information revealed substantial variation in DDPM-generated samples across runs, although SC patterns were generally more apparent in HCP than in NIMH (Figure S2). The residual matrices revealed substantially smaller absolute differences in HCP than in NIMH when visualized using a common color scale (Figure 4). These differences should not be interpreted as evidence of superior generative performance on HCP, as the edge weights in HCP were smaller than those in NIMH. When the color scale was adjusted to the HCP dataset, residual patterns became visible for DDPM.

**Figure 4.**
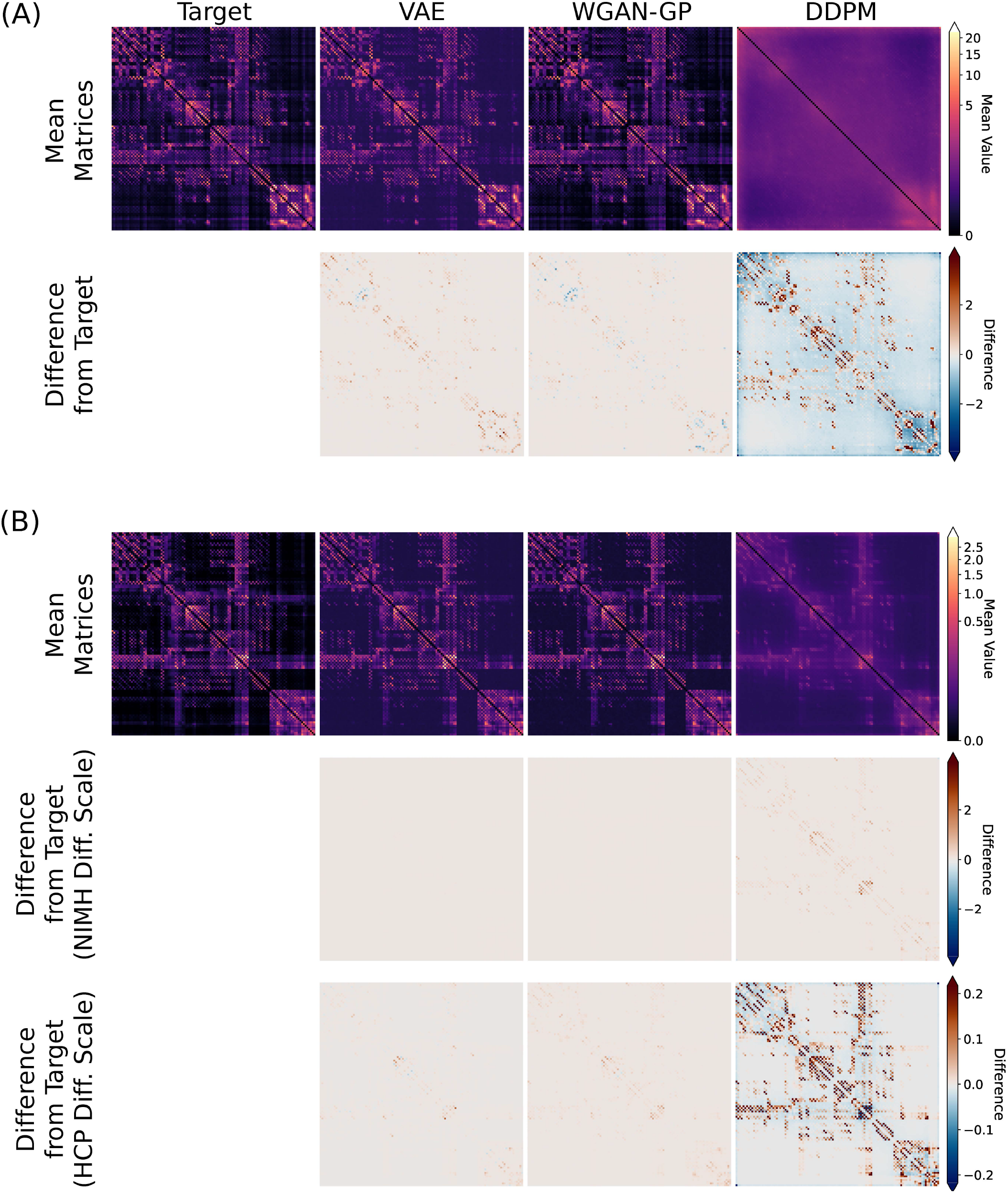
Qualitative and visual comparison of adjacency matrices for the SC datasets. (A) NIMH and (B) HCP datasets. Top row: Mean adjacency matrices of the test set and the generated graphs. For each model, the mean matrix was calculated by pooling all generated samples across runs. Power-law color normalization (*γ* = 0. 3) was applied to enhance edge visibility. Subsequent rows: Corresponding residual matrices (test - generated). The 1^st^ and GG^th^ percentiles were calculated from the residuals pooled across all models, and the larger absolute value was used to set symmetric color limits. In (A), the residuals are displayed using the NIMH-specific color limits. In (B), the second row uses the same color limit as NIMH, while the third row uses HCP-specific color limits. *Abbreviations*: VAE = variational autoencoder, WGANGP = Wasserstein GAN with gradient penalty, DDPM = denoising diffusion probabilistic models, SC = structural connectivity, NIMH = NIMH Healthy Volunteer Dataset, HCP = Human Connectome Project Young Adult S1200 dataset.

In the MMD-based evaluation, WGAN-GP achieved the lowest MMD values for most metrics and thresholding criteria in both datasets. VAE generally performed better than DDPM in NIMH; however, their relative performance varied by metric (Figure 5). For modularity and small-worldness, WGAN-GP achieved the smallest Wasserstein distances, followed by VAE, whereas DDPM-generated distributions deviated the most from the test distributions (Figure 5B, C). The scatter plots also revealed considerable variation in DDPM performance across runs.

**Figure 5.**
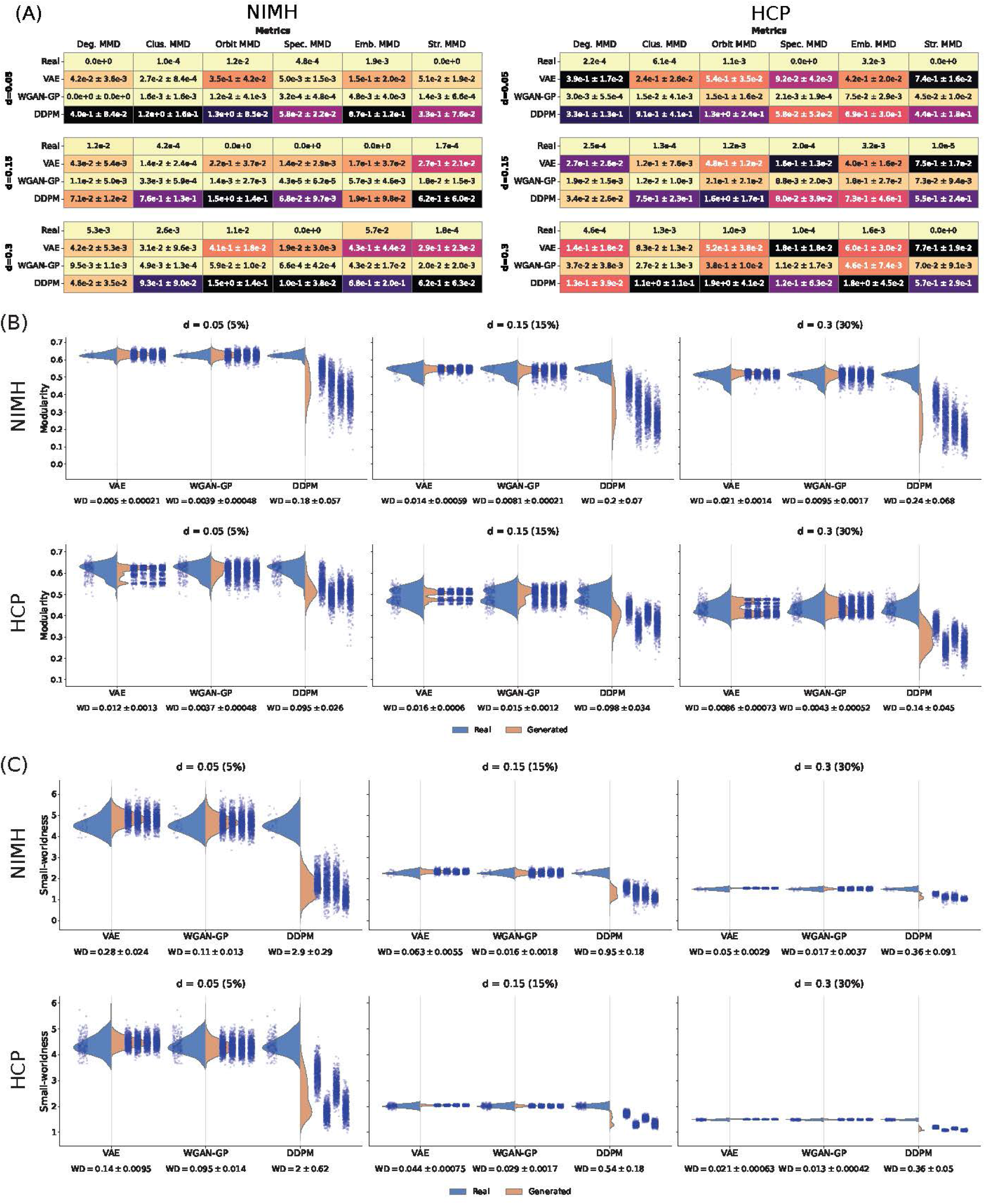
Quantitative evaluation results of generative models across SC datasets. (A) Heatmap of the MMD results for the SC dataset for each thresholding criterion. Cells are colored based on each metric’s value across all models and thresholding criteria, with lighter colors indicating better performance. (B) Distributions of modularity scores and Wasserstein distances between the generated graphs and the test set. (C) Distributions of small-worldness and Wasserstein distances between generated graphs and the test set. For all panels, edge density (*d*) indicates the proportion of strongest edges retained in each graph. In (B) and (C), violin plots show the distributions pooled across four independent runs, with scatter points representing individual runs, arranged from left to right in the order of runs 1-4. MMD and Wasserstein distance values are reported as mean ± standard deviation across four independent runs. *Abbreviations*: VAE = variational autoencoder, WGAN-GP = Wasserstein GAN with gradient penalty, DDPM = denoising diffusion probabilistic models, SC = structural connectivity, NIMH = NIMH Healthy Volunteer Dataset, HCP = Human Connectome Project Young Adult S1200 dataset.

The individual-variability analyses further differentiated the models (Figure 6). WGAN-GP most closely reproduced the edge-wise variability of the test data in both datasets, whereas VAE generally showed reduced variability and DDPM exhibited larger deviations across edges. Similarly, the distributions of nearest-neighbor and pairwise distances for WGAN-GP were closest to those of the test data, while VAE produced smaller distances. DDPM produced larger and more widely distributed distances, with substantial variation across runs.

**Figure 6.**
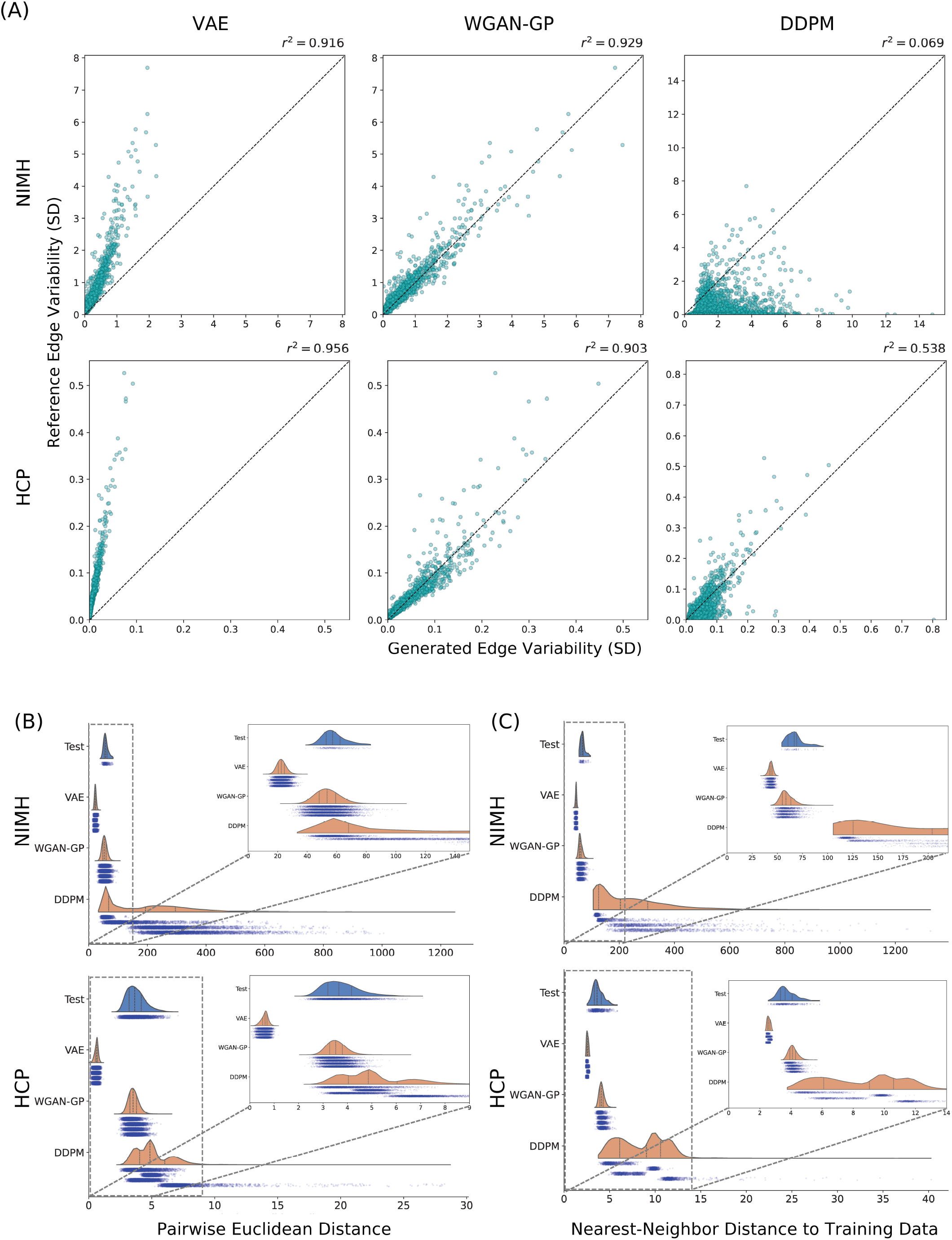
Diversity and novelty analyses of generated SC data in the NIMH and HCP datasets. (A)Edge-wise variability analysis. Each point represents an edge, comparing the standard deviation of its weight across the generated graphs (x-axis) and the test graphs (y-axis). Edge-wise standard deviations were calculated separately for each run and then averaged across runs. The diagonal line indicates equal variability between the test and generated data. The *r*^2^ values indicate the squared Pearson’s correlation coefficient. (B) Pairwise distance analysis. The violin plot shows the distributions of Euclidean distances between all pairs of samples within the test or generated samples, pooled across runs. (C) Nearest-neighbor analysis. The violin plot shows the distributions of Euclidean distances from each test or generated sample to its nearest sample in the corresponding training dataset, pooled across runs. In both (B) and (C), the scatter plots below the violin plots show the distributions of individual runs, arranged from top to bottom in the order of runs 1-4. *Abbreviations*: VAE = variational autoencoder, WGAN-GP = Wasserstein GAN with gradient penalty, DDPM = denoising diffusion probabilistic models, SC = structural connectivity, NIMH = NIMH Healthy Volunteer Dataset, HCP = Human Connectome Project Young Adult S1200 dataset.

**Table 1.**
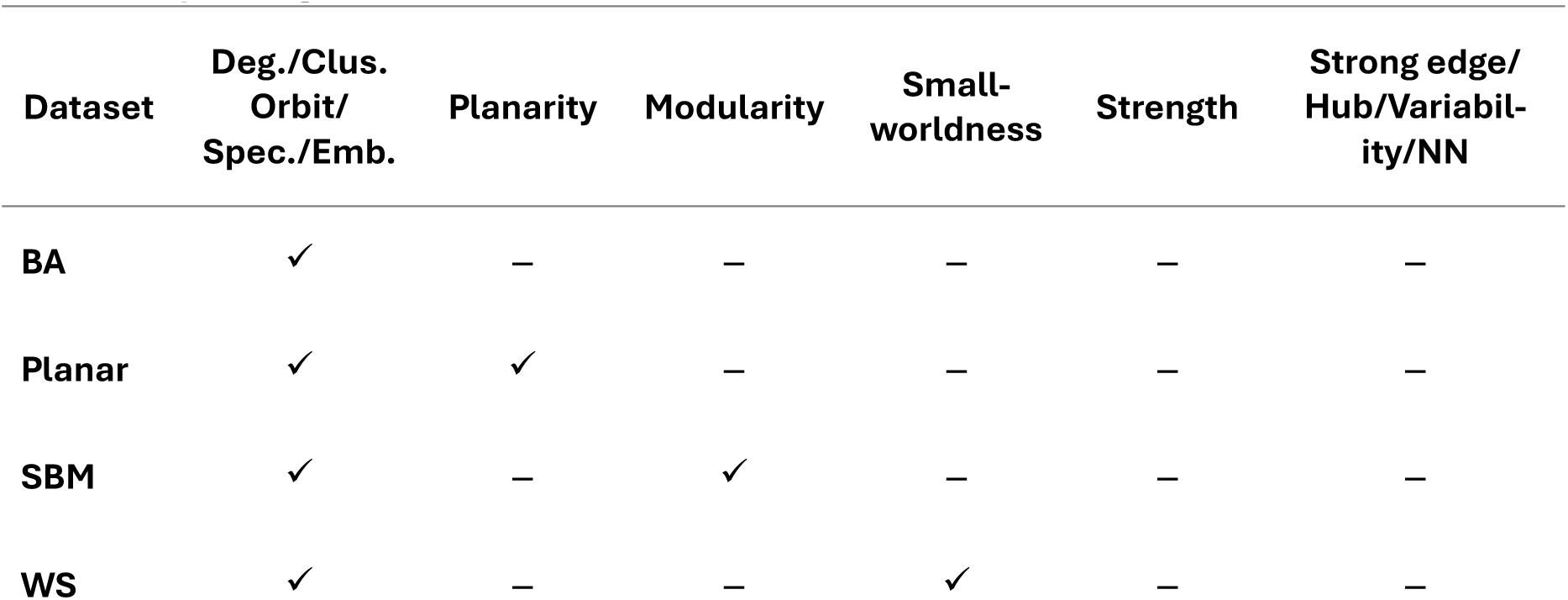
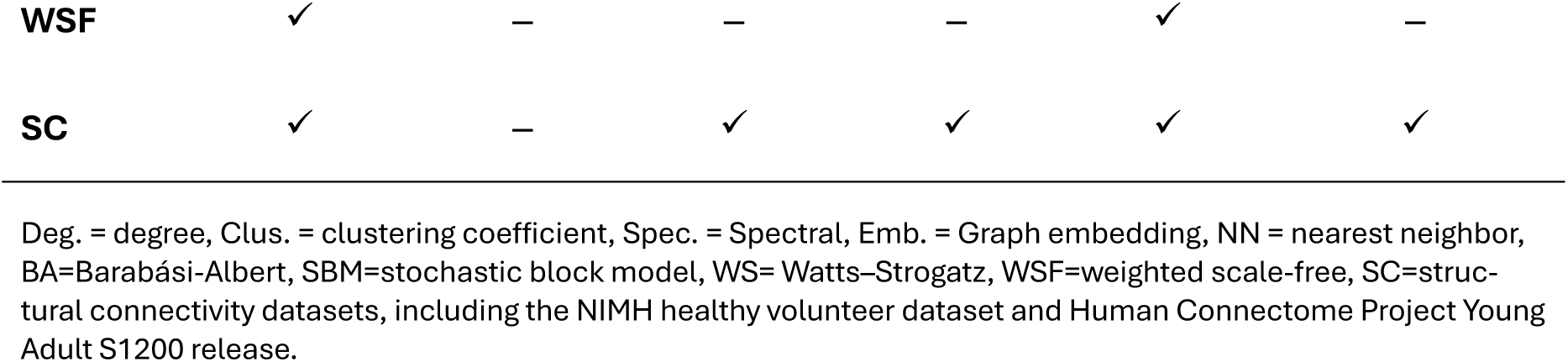
Summary of evaluation metrics used for each dataset. A check mark indicates that the corresponding metric was used for that dataset.

| Dataset | Deg./Clus. Orbit/Spec./Emb. | Planarity | Modularity | Small-worldness | Strength | Strong edge/Hub/Variabil-ity/NN |
|---|---|---|---|---|---|---|
| **BA** | ✓ | – | – | – | – | – |
| **Planar** | ✓ | ✓ | – | – | – | – |
| **SBM** | ✓ | – | ✓ | – | – | – |
| **WS** | ✓ | – | – | ✓ | – | – |
| **WSF** | ✓ | – | – | – | ✓ | – |
| **SC** | ✓ | – | ✓ | ✓ | ✓ | ✓ |

Deg. = degree, Clus. = clustering coefficient, Spec. = Spectral, Emb. = Graph embedding, NN = nearest neighbor, BA=Barabási-Albert, SBM=stochastic block model, WS= Watts–Strogatz, WSF=weighted scale-free, SC=struc-tural connectivity datasets, including the NIMH healthy volunteer dataset and Human Connectome Project Young Adult S1200 release.

VAE and WGAN-GP also performed well at reconstructing important connections and hub nodes in the SC data (Table 2). VAE and WGAN-GP achieved high Jaccard indices and Spearman correlation coefficients in both datasets, indicating substantial strong edge overlap and high agreement in node-strength rankings. DDPM showed lower Jaccard indices and node-strength correlations than the other models, although both measures improved in HCP. The overlap and differences in strong edges across individual runs are visualized in the Supplementary Information. The results for each individual run, including MMD, Wasserstein distance, Jaccard index, and hub preservation metrics, are provided in Supplementary Tables S9-S13.

**Table 2.** Strong-edge and hub preservation in the NIMH and HCP datasets. Jaccard indices (top 1%, 5%, and 10% of edges) and Spearman’s *ρ* are reported as mean ± SD across runs. Sign ificant correlations are indicated as the number of significant runs/total runs (*p* < 0. 05).

| Dataset | Model | Strong edge preservation |  |  | Hub preservation |  |
|---|---|---|---|---|---|---|
|  |  | Jaccard 1% | Jaccard 5% | Jaccard 10% | Spearman *ρ* | Significant runs |
| NIMH | VAE | 0.8593±0.0214 | 0.8921±0.0044 | 0.9084±0.0022 | 0.9904±0.0015 | 4/4 |
|  | WGAN-GP | 0.8791±0.0133 | 0.8854±0.0067 | 0.9050±0.0056 | 0.9908±0.0013 | 4/4 |
|  | DDPM | 0.0739±0.0648 | 0.1359±0.0701 | 0.1940±0.0685 | 0.2693±0.1227 | 3/4 |
| HCP | VAE | 0.9704±0.0240 | 0.9836±0.0030 | 0.9844±0.0028 | 0.9988±0.0002 | 4/4 |
|  | WGAN-GP | 0.9776±0.0149 | 0.9763±0.0048 | 0.9814±0.0044 | 0.9988±0.0002 | 4/4 |
|  | DDPM | 0.3539±0.3142 | 0.3920±0.2645 | 0.4087±0.1991 | 0.7121±0.2391 | 4/4 |

VAE=variational autoencoder, WGAN-GP=Wasserstein GAN with gradient penalty, DDPM=denoising diffusion probabilistic models, NIMH = NIMH Healthy Volunteer Dataset, HCP =Human Connectome Project Young Adult S1200 release.

## 4. Discussion

In this study, we compared the generative characteristics of three deep generative models using five synthetic graph datasets with different structural properties and two empirical SC datasets. Evaluation of the synthetic datasets revealed shared limitations in reproducing complex topological properties, as well as distinct model-specific generative biases. VAE tended to overemphasize common structural patterns, WGAN-GP showed a balanced ability to reproduce structures, and DDPM captured stochastic connection patterns but struggled to reproduce coordinate-specific patterns and showed substantial performance variation across runs. Evaluation on the two empirical SC datasets further revealed corresponding differences in model performance, where WGAN-GP most closely reproduced both the topological properties and inter-subject variability of the test data, while VAE underestimated sample variability and DDPM showed variation across runs.

### 4.1. Model-Specific Characteristics Revealed by Synthetic Graphs

The synthetic datasets were used not to directly determine the best model for SC data, but to examine how generative models reproduce different graph properties and to identify model-specific generative biases. Here, we discuss the shared limitations of the evaluated models in reproducing complex graph properties, followed by their distinct generative characteristics and potential explanations related to network architecture and training objectives.

The difficulty of reproducing strict global constraints, which require macroscopic consistency in edge presence across the entire network, suggests that the evaluated vector- and image-based generative models rely on optimizing statistical resemblance and may struggle to implicitly learn the complex topological rules that govern these properties. All models struggled to capture the degree distribution of BA (Figure 3A) and the validity of Planar (VAE: 0.15, WGAN-GP: 0.0, DDPM: 0.0; Figure 3A). The small sample size of the Planar dataset may also have contributed to the low validity of the generated graphs. Similarly, none of the models fully reproduced the small-world structure of the WS dataset, with generated graphs consistently showing lower small-worldness values than the target graphs. Although WGAN-GP and DDPM visually generated some off-diagonal edges, their lower small-worldness values suggest that generating visually plausible long-range edges alone was insufficient to reproduce the global organization of edges needed to balance local clustering and global connectivity. These tendencies were consistent across runs, suggesting that they reflect generative biases rather than isolated variation between runs. These results highlight the limitations of the evaluated implementations in capturing higher-order dependencies among edges, suggesting that explicitly incorporating structural constraints may be beneficial.

The transition to weighted graphs also revealed a shared limitation across all models. Although WGAN-GP and DDPM achieved relatively low MMD values for WSF, all architectures tended to underestimate edge weights, as shown in the mean adjacency matrices (Figure 2E). This suggests that even when topological features are reproduced to some extent, accurately reproducing continuous edge weights remains a challenge for these generative models. Together with the aforementioned difficulties in reproducing global structural constraints, these findings highlight the need to carefully select a baseline based on its specific generative biases rather than expecting perfect reproduction.

The VAE consistently exhibited a strong bias towards generating common edges, struggling to capture rare structural connections and global constraints. This intrinsic bias was clearly shown in the results. Although VAE achieved the highest scores on 4 of 6 metrics for the Planar dataset (Figure 3A), visual inspection shows an exaggerated concentration of edges around the diagonal and visually implausible graphs (Figure 2B). This same overemphasis led to an inflated modularity of nearly 0.5 in the SBM dataset, due to excessive intra-community connections and fewer inter-community connections, resulting in a loss of crucial long-range shortcuts in WS. For WSF, the VAE generated mean-like outputs that lacked sparse edge patterns, resulting in poor performance in all metrics (Figure 3A, Figure S1E). Thus, while this over-smoothing could deceptively lower MMD values for common features, it fundamentally hinders strict topological generation. This tendency was consistently observed across multiple independent runs, suggesting that it is a reproducible model-specific bias rather than an artifact of a particular run. This phenomenon is a well-documented characteristic of VAEs; optimizing the element-wise loss forces the model to predict the average of multiple data points mapped to the same latent representation, leading to over-smoothed outputs in which rare or sharp features are lost.^36,37^

DDPM showed a distinct advantage in capturing stochastic, texture-like connection patterns, but its architecture and diffusion process may have contributed to difficulty reproducing coordinate-specific graph structures. Its performance in SBM, together with the blurred mean adjacency matrix (Figure 2C), may reflect its capacity to capture the underlying connection probabilities rather than overfitting to rigid edges, consistent with previous findings on diffusion-based texture synthesis.^38^ Although DDPM successfully reproduced rigid, coordinate-specific graph patterns in some runs, the quality of their reproduction varied substantially across runs, particularly in the Planar and WSF datasets (Figure S1B, E). Given that run 1 successfully reproduced these patterns, the observed instability may reflect sensitivity to training conditions. This discrepancy could partly reflect the inductive bias of the CNN architecture used in DDPM. MLPs process the entire input simultaneously through fully connected layers, which allows them to learn global relationships between distant elements of the adjacency matrix. This aligns with findings in Tabular data generation tasks, where MLP-based models are reported to outperform CNN-based models,^39^ suggesting that MLPs are better suited for generating data where capturing global relationships is more crucial than local features. In contrast, CNNs employ shared convolutional filters, introducing translation equivariance that is advantageous for natural images, but may be less suitable for adjacency matrices where each position represents a connection between specific nodes. Furthermore, previous research has pointed out that adding Gaussian noise to adjacency matrices during the forward diffusion process destroys their sparsity and graph-theoretic properties. ^40^ This disruption of structural information may have contributed to the difficulties in reliably reproducing coordinate-specific edge patterns observed in our experiments. Since DDPM is the only model that adopts a CNN-based architecture, the observed limitations of DDPM should be interpreted as those of the specific U-Net-based implementation evaluated here, rather than as general limitations of diffusion models.

Overall, WGAN-GP demonstrated highly balanced generative performance across the synthetic datasets, which may be attributed to its adversarial training framework. Specifically, the model generally performed among the top two models across the evaluated metrics in multiple runs, with only limited cases of lower rankings, while achieving the best performance for modularity and small-worldness. It also avoided the over-smoothing or fixed-coordinate collapse observed in VAE and DDPM. However, it is not without limitations: in addition to the universal challenges, visual inspection reveals that continuous structures were fragmented in the Planar and WS datasets (Figure S1B, D), and the model did not achieve quantitative results comparable to the test dataset. This suggests that adversarial training, rather than element-wise reconstruction losses, may contribute to better global distributional matching, although it does not necessarily ensure the reproduction of strict local topological rules.

### 4.2. Model-Specific Biases in Empirical SC Matrix Generation

The SC results were interpreted in light of the model-specific biases identified in the synthetic datasets. By extending the comparison from controlled synthetic graphs to empirical SC datasets, we examined whether the generative biases identified for individual graph properties were also observed in brain networks. Across the two SC datasets, WGAN-GP consistently demonstrated the best overall performance across the evaluated metrics. However, the relative performance of VAE and DDPM differed between the datasets. While VAE clearly outperformed DDPM on the NIMH dataset, the performance gap narrowed on the HCP dataset, particularly in the MMD-based evaluations. This section discusses how the ability to preserve coordinate-specific patterns, common structural backbones, stochastic variability, and weighted edges translated into empirical SC generation.

Consistent with the findings from synthetic datasets discussed in Section 4.1, VAE and WGAN-GP also demonstrated a high capacity to preserve fixed-coordinate structural patterns in empirical SC data. The strong pattern retention is visually plausible (Figure 4) and quantitatively supported by high preservation scores for both strong edges and hubs. Across both the NIMH and HCP datasets, VAE and WGAN-GP achieved Jaccard indices ranging from 0.86 to 0.98 across all thresholds and a near-perfect Spearman correlation coefficient of 0.990-1.00 for node strength preservation (Table 2). These results suggest that the evaluated MLP-based architectures may contribute to preserving fixed-coordinate structural patterns in SC matrices, which may reflect the stable organization of major anatomical connections in the brain.

Conversely, as anticipated from the architectural bias discussed in Section 4.1, DDPM struggled to pinpoint exact edge coordinates in the NIMH dataset. Qualitative observations reveal that DDPM generated SC-like connection textures but failed to consistently reproduce the prominent connectivity patterns in the reference dataset across individual samples. This lack of coordinate consistency resulted in a blurry mean adjacency matrix and high MMD scores across most metric-threshold combinations (Figure 4A). In the HCP dataset, DDPM reproduced SC patterns more clearly than in the NIMH dataset (Figure 4B), although its performance varied considerably across runs, consistent with synthetic datasets. However, when visualized using a dataset-specific color scale, DDPM exhibited larger residuals than VAE and WGAN-GP. This improvement suggests that the difficulty of reproducing fixed-coordinate patterns may partly depend on sample size or other dataset characteristics. Nevertheless, even with the larger HCP dataset, DDPM’s overall performance did not reach that of the best-performing model. These contrasting results underscore that preserving coordinate-specific connectivity patterns is important for accurately reproducing empirical SC data.

Although VAE and WGAN-GP successfully reproduced the average spatial patterns of the SC matrices in both datasets, population-level distributional fidelity revealed further differences between the models. WGAN-GP achieved the lowest MMD values among compared models across most graph-theoretical features. However, the extent of this fidelity varied between datasets. In the NIMH dataset, WGAN-GP achieved MMD values comparable to the reference baseline across most metrics, particularly for the top 5% of the edges, whereas larger discrepancies remained in the HCP dataset (Figure 5A). This superiority was also evident in modularity and small-worldness, where WGAN-GP consistently achieved the smallest Wasserstein distances, except for a single run in the modularity evaluation (Figure 5B, C). Notably, although DDPM generally struggled on the NIMH dataset, it achieved lower MMD values than VAE for degree, spectral, and node strength in the HCP dataset (Figure 5A). Nevertheless, this improvement did not consistently extend to modularity and small-worldness, where DDPM showed larger deviations from the test distributions. These findings suggest that accurately reproducing average connectivity patterns does not necessarily guarantee faithful reproduction of graph-theoretical feature distributions, and that reproducing individual graph statistics does not necessarily preserve broader topological properties.

Interestingly, while VAE and WGAN-GP struggled to replicate global topological properties in synthetic datasets, they demonstrated a remarkable capability to approximate these structures in empirical SC data. Specifically, WGAN-GP successfully reproduced the distribution peaks of modularity and small-worldness, as well as their gradual shifts across varying sparsity thresholds, in both SC datasets (Figure 5). VAE also reproduced the small-worldness distributions in both datasets and modularity distributions in NIMH relatively well. This contrast between synthetic and empirical datasets could be attributed to the inherent underlying rules of the SC datasets. Unlike purely mathematical graphs, SC networks are constrained by human brain anatomy, formed through optimization between wiring costs and network efficiency.^42^ Models can capitalize on these probability biases of edge presence (e.g., highly probable anatomically anchored long-range connections). By learning these spatial probability biases, the models could indirectly approximate modularity and small-worldness. Consequently, the successful reproduction of these graph-theoretical properties in the SC datasets could be better explained by anatomy-assisted approximation than by directly learning underlying topological rules.

Individual-level evaluations revealed distinct model-specific characteristics in reproducing inter-subject variability (Figure 6). VAE exhibited the lowest edge-wise variability and smaller pairwise distances across both SC datasets. Furthermore, its nearest-neighbor distances to the training data were smaller than those of real test samples. These findings are consistent with the over-smoothing tendency observed in synthetic datasets. Although VAE captured the relative patterns of edge-wise variability, as evidenced by high Pearson correlations, its inferior performance to WGAN-GP even in MMD metrics based on scale-independent binary graph properties after percentile thresholding indicates that its limitations extend beyond the absolute scale of edge weights to the reproduction of topological properties across individual graphs. DDPM, in contrast, showed greater variability than the real data across all three individual-level evaluations. Considering its discrepancies from the real data in the MMD evaluation, this diversity may reflect deviations from realistic samples rather than faithful reproduction of inter-subject variability. WGAN-GP yielded results closest to the real data across all three evaluations. These findings further support the balanced performance of WGAN-GP, demonstrating its ability to reproduce realistic inter-subject variability in addition to the graph-theoretical feature distributions evaluated by MMD. Nevertheless, none of the models showed clear signs of severe mode collapse or exact memorization of training samples, such as a concentration of pairwise or nearest-neighbor distances near zero.

These results suggest that WGAN-GP is a promising candidate among the evaluated models for future studies on SC matrix generation. Since data augmentation aims to increase data diversity for better generalization, the over-smoothing tendency of VAEs, biasing generation toward high-probability edges, could limit its effectiveness for data augmentation in classification or regression tasks. Similarly, DDPM struggled to reliably preserve the exact sparse coordinates of the connectivity patterns. Conversely, WGAN-GP captured overall network traits, especially for strong edges, preserving graph variety. WGAN-GP not only outperformed other models in maintaining data diversity (Figure 6) but also showed balanced performance in structural fidelity, achieving the best performance on most metrics across datasets and runs. Moreover, it reproduced major connections and hubs at the group level (Table 2). Preserving these major structural hubs is crucial, as neurological disorders, such as AD and frontotemporal dementia, selectively target these hubs.^43^ This suggests a possible direction for future research into structural connectivity abnormalities as biomarkers. However, the effectiveness of data augmentation using these generative models in downstream tasks, such as prediction or classification, remains to be validated.

### 4.3. Limitations and Future Directions

Several limitations should be acknowledged. First, the observed performance differences may depend on the specific implementations, architectures, and hyperparameters used in this study. In particular, DDPM was implemented with a U-Net architecture, whereas VAE and WGAN-GP were implemented with MLP-based architectures. Therefore, the effects of the generative paradigm and network architecture cannot be fully separated. In addition, the use of different criteria for hyperparameter optimization and early stopping across models may have influenced model selection and the resulting performance comparisons. Although we performed hyperparameter optimization for DDPM, we did not explore alternative noise schedules or numbers of diffusion steps, which may have affected its performance. This study focused on three representative deep generative models and did not include comparisons with non-deep generation methods, such as SBM, edge-wise resampling, or other statistical graph models. Graph-specific generative models, such as VGAE,^44^ GraphVAE,^45^ NetGAN,^46^ SPECTRE,^1^ and DiGress,^40^ employ approaches that explicitly utilize graph characteristics and could be alternative approaches for addressing higher-order constraints. However, their application to SC data with anatomical node correspondence and weighted edges needs further exploration. Our findings should be interpreted as a comparison among the evaluated implementations rather than as conclusions applicable to all graph generation methods. Future studies should evaluate alternative architectures, including MLP-based diffusion models or graph-specific architectures, to determine whether the observed tendencies generalize beyond the present implementation.

Second, our preprocessing and evaluation procedures pose some limitations. Our decision not to apply additional scaling to edge weights before training might have affected training stability and the optimal performance of the generative models. Additionally, the empirical SC matrices evaluated in this study were derived from dMRI tractography, which is subject to false-positive/false-negative connections, distance bias, and dependence on preprocessing and tractography methods.^47^ Furthermore, results in this paper are confined to generating graph adjacency matrices with a fixed node order. Applying node ordering could introduce dataset-specific spatial patterns that may have affected model learning and performance. For example, the community-based ordering for SBM and k-core ordering for WS may have emphasized community structures and topological patterns, respectively. Therefore, our results cannot be generalized to permutation-invariant graph generation. For synthetic datasets, we thresholded the generated graphs to match the edge densities of the original graphs. Consequently, the evaluation does not directly assess the models’ ability to reproduce graph sparsity. Instead, the results reflect their ability to reproduce structural properties after edge-density adjustment.

Finally, this study focused on the structural fidelity of generated graphs rather than directly evaluating their impact on downstream prediction or classification tasks. Although the present results suggest that WGAN-GP provides a balanced baseline for preserving major SC structures, its practical effectiveness as a data augmentation method should be validated in task-specific settings.

## 5. Conclusion

In this study, we systematically compared three representative deep generative models—VAE, WGAN-GP, and DDPM—to elucidate their model-specific generative biases and capabilities in capturing the topological features of ROI-level weighted SC matrices defined on a fixed anatomical parcellation. By combining synthetic graph datasets with controlled characteristics and empirical SC datasets, we examined not only overall generation quality but also which graph properties each model reproduced well or poorly, as well as how well the models reproduced inter-subject variability and sample diversity in the SC datasets. Among the evaluated models, WGAN-GP emerged as the most balanced baseline in terms of structural fidelity, showing balanced performance across the synthetic datasets and achieving the best results on most evaluation metrics in both SC datasets. VAE and WGAN-GP both preserved prominent connections and hub structures; however, WGAN-GP more closely reproduced inter-subject variability and sample diversity.

At the same time, our findings highlight the limitations of directly applying standard image- or vector-based generative models to brain graph data. All evaluated models struggled to fully reproduce complex graph-theoretic properties, such as higher-order edge dependencies underlying planarity in the Planar dataset, degree distributions in the BA dataset, and edge weights in the WSF dataset. These results support approaches that explicitly account for graph structure by incorporating graph-specific structural constraints or biologically meaningful priors into the training or generation process to improve the fidelity of SC matrix generation. Overall, this empirical comparison provides insights into the models’ capabilities and limitations for various synthetic graphs and empirical SC matrices, offering an empirical basis for future research on SC generation.

## Acknowledgement

This work was supported by the Japan Society for the Promotion of Science KAKENHI Grant Number JP24K03028, and in part by grants-in-aid from the Harris Science Research Institute of Doshisha University. We acknowledge the significant contributions of the NIMH Healthy Volunteer Dataset, which provided the MRI data and task scores. Data were provided in part by the Human Connectome Project, WU-Minn Consortium (Principal Investigators: David Van Essen and Kamil Ugurbil; 1U54MH091657) funded by the 16 NIH Institutes and Centers that support the NIH Blueprint for Neuroscience Research; and by the McDonnell Center for Systems Neuroscience at Washington University.

## Data Availability Statement

The four synthetic datasets were generated using established algorithms (Barabási-Albert model, stochastic block model, Watts–Strogatz model, weighted scale-free model). The source code for this study and the generated synthetic datasets are available on GitHub (https://github.com/MIS-Lab-Doshisha/gensc-bench). The planar dataset is available under the MIT license (https://github.com/KarolisMart/SPECTRE).^1^ The structural connectivity dataset was derived from the NIMH Healthy Volunteer Dataset^2^ and is available in the OpenNeuro repository (https://doi.org/10.18112/openneuro.ds004215.v1.0.0) under a Creative Commons Zero (CC0) license. The preprocessed MRI data from the Human Connectome Project Young Adult S1200 release are available through Amazon S3 (s3://hcp-openaccess/HCP_1200) to registered users, subject to the HCP Open Access Data User Terms.

## Funding Statement

This work was supported by the Japan Society for the Promotion of Science KAKENHI (ROR: https://ror.org/00hhkn466) Grant Number JP24K03028, and the Harris Science Research Institute of Doshisha University.

## Conflict of Interest Disclosure

The authors declare no conflicts of interest.

## Ethics Approval Statement

This study used publicly available datasets and algorithmically generated synthetic data. Therefore, ethical approval was not required.

## Patient consent statement

Not applicable.

## Permission to reproduce material from other sources

Not applicable.

## Clinical trial registration

Not applicable.

## Notes

https://github.com/MIS-Lab-Doshisha/gensc-bench

### Competing Interest Statement

The authors have declared no competing interest.

### Summary of Updates

The major revisions include: 1.Re-evaluation of all experiments: We corrected issues in the WGAN-GP training procedure and random seed initialization, and reran all experiments, including hyperparameter optimization. 2.Expanded empirical evaluation: We added the Human Connectome Project Young Adult dataset (N = 1,065) to assess the consistency of our findings across independent structural connectivity datasets. 3.Assessment of run-to-run variability: We conducted four training runs with different training/validation splits and random seeds to evaluate the variability in generative performance. 4.Additional diversity and novelty analyses: We introduced edge-wise variability, pairwise distance, and nearest-neighbor analyses to assess the diversity and potential memorization of generated samples. 5.Clarification of the study's scope and limitations: We refined the manuscript to emphasize its contribution as a systematic empirical benchmark and to distinguish structural fidelity from effectiveness in downstream data augmentation tasks.

