## Supplementary for "Comparative Evaluation of Deep Generative Models for Capturing Topological Features in Brain Structural Connectivity Matrices"

### Supplementary Information

#### S.1. Model Details

##### S.1.1. Variational Autoencoder (VAE)

The loss function is defined as follows:

$$\mathcal{L} = -\mathbb{E}_{q_\phi(z|x)}[\log p_\theta(x|z)] + D_{\mathrm{KL}}\big(q_\phi(z|x)|p(z)\big)$$

where $q_\phi(z|x)$ is the output of the encoder, $p_\theta(x|z)$ is the output of the decoder, and $p(z)$ is the prior distribution over the latent space, which was set to a standard normal distribution in this study.

The encoder consisted of repeated blocks, each comprising a linear layer, a batch normalization layer, and a nonlinear activation function. For the synthetic dataset, LeakyReLU was used as the activation function, whereas ReLU was employed for the SC dataset. The decoder mirrored the architecture of the encoder, with the final layer being a Softplus activation function to ensure non-negative outputs. For the synthetic datasets, the number of layers was fixed to two for both the encoder and the decoder. For the SC dataset, the number of layers was treated as a hyperparameter and optimized during training.

##### S.1.2. Wasserstein GAN with Gradient Penalty

The loss functions for the critic $C$ and the generator $G$ are defined as follows:

$$\mathcal{L}_C = \mathbb{E}_{\tilde{x}\sim p_g}[C(\tilde{x})] - \mathbb{E}_{x\sim p_r}[C(x)] + \lambda\mathbb{E}_{\hat{x}\sim p_{\hat{x}}}[(\| \nabla_{\hat{x}} c(\hat{x}) \|_2 - 1)^2]$$
$$\mathcal{L}_G = -\mathbb{E}_{\tilde{x}\sim p_g}[C(\tilde{x})]$$

where $p_g$ is the distribution of generated data, $p_r$ is the distribution of real data, and $p_{\hat{x}}$ is the distribution of samples interpolated between real and generated data. The third term in the critic loss is the gradient penalty, with $\lambda$ being the penalty coefficient.

The generator architecture consisted of repeated blocks, each comprising a linear layer, a batch normalization layer, and a nonlinear activation function. For the synthetic dataset, LeakyReLU was used as the activation function, whereas ReLU was employed for the SC dataset. The critic mirrored the architecture of the generator, except that it included dropout layers instead of batch normalization layers. For the synthetic datasets, the number of layers was fixed to two for both the generator and the critic. For the SC dataset, the number of layers was treated as a hyperparameter and optimized during training.

During training, the critic was updated three times for each generator update ($n_{\mathrm{critic}} = 3$), using the same mini-batch of real samples for all three critic updates. The gradient penalty coefficient was fixed at $\lambda = 10$.

##### S.1.3. Denoising Diffusion Probabilistic Models (DDPM)

The loss function for training the model is defined as the mean squared error between the added noise and the predicted noise:

$$\mathcal{L} = \mathbb{E}_{x_0,\epsilon,t}[\| \epsilon - \epsilon_\theta(x_t, t) \|^2]$$

where $x_0$ is the original data, $\epsilon$ is the added noise, $x_t$ is the noisy data at time step $t$, and $\epsilon_\theta(x_t, t)$ is the output of the neural network predicting the noise. The noise schedule followed the original DDPM implementation by Ho et al., with a linear β schedule.

For the synthetic datasets, the base channel number of the U-Net was fixed to 16. For the SC dataset, the base channel number was treated as a hyperparameter and optimized during training.

### S.2. Dataset Details

#### S.2.1. Synthetic Dataset Generation

The Barabási-Albert (BA), Stochastic Block Model (SBM), and Watts-Strogatz (WS) datasets were generated using the NetworkX[1] library. The specific parameters used for generating each dataset are as follows:

**Barabási-Albert (BA):** Each graph was constructed with an initial connected network of 5 nodes, and each new node connecting to 4 existing nodes.

**Planar:** The dataset consists of 200 planar graphs generated using Delaunay triangulation and obtained from the dataset provided by Martinkus et al.[2]

**Stochastic Block Model (SBM):** Graphs were generated with two communities, each containing 28 to 36 nodes. The intra-community connection probability was set to 0.3, and the inter-community connection probability was set to 0.05.

**Watts-Strogatz (WS):** Each node was initially connected to its six nearest neighbors, with a rewiring probability of 0.05.

**Weighted Scale-Free (WSF):** Each graph was constructed with an initial connected network of 9 nodes, and each new node connecting to 8 existing nodes.

#### S.2.2. Structural Connectivity Reconstruction

SC matrices were constructed using closely matched procedures. Tissue-specific response functions were estimated using the Dhollander algorithm,[3] followed by fiber orientation distribution estimation with constrained spherical deconvolution (CSD) and multi-tissue intensity normalization. Anatomically constrained probabilistic tractography was performed using the iFOD2 algorithm[4] with 10 million streamlines, followed by SIFT2.[5] The whole brain was parcellated into 116 regions based on the automated anatomical labeling (AAL) atlas.[6] SC matrices were constructed by summing the SIFT2 weights of streamlines connecting each pair of regions, with edge weights additionally scaled by the inverse node volumes.

In the NIMH dataset, the MRI data were preprocessed using QSIprep (version 0.15.4),[7] as previously described,[8] followed by SC reconstruction including fiber orientation estimation using single-shell three-tissue CSD (SS3T-CSD).[9]

In the HCP dataset, SC matrices were reconstructed from the preprocessed diffusion- weighted images and T1-weighted images provided by HCP project using MRTrix3 (version 3.0.8).[10] Unlike the NIMH dataset, fiber orientation distributions were estimated using multi-shell multi-tissue CSD (MSMT-CSD).[11]

#### S.3. Dataset Separation Details

For the synthetic datasets and the HCP dataset, samples were randomly divided into the discovery and test sets. The discovery set was then randomly partitioned into four folds of approximately equal size.

For the NIMH dataset, a different splitting procedure was employed due to its relatively small sample size. Participants were first sorted by the age-adjusted fluid intelligence scores from the NIH Toolbox Cognition Battery.[8,12] Every fourth participant in the sorted list (1st, 5th, 9th, ..., and 105th) was assigned to the test set, while the remaining participants were assigned to the discovery set.

The discovery set was then divided into four folds using the same rank-based assignment procedure. Participants in the discovery set were assigned to four folds in a round-robin manner according to their fluid intelligence score. Specifically, participants ranked 1st, 5th, 9th, and so on were assigned to fold 1; those ranked 2nd, 6th, 10th, and so on were assigned to fold 2; and the same procedure was applied to folds 3 and 4.

#### S.4. Computational Environment

All experiments were executed on a server with an NVIDIA RTX 4080 Super GPU, an Intel Core i9-14900k CPU, and 32GB of RAM.

#### S.5. Training Configurations and Selected Hyperparameters

The training configurations and hyperparameter search spaces for the synthetic dataset, the NIMH dataset, and the HCP dataset are summarized in Table S1. For the synthetic dataset, the model capacity was constrained for a controlled comparison. The search space for the VAE latent dimension was intentionally kept smaller than that of WGAN-GP to prevent posterior collapse.

In contrast, architecture-related hyperparameters were treated as tunable for the Structural Connectivity (SC) dataset. This approach allows the models to be compared under dataset-specific optimized settings. The final selected hyperparameters are summarized in Table S2 and Table S3.

Each hyperparameter optimization for both synthetic and SC datasets consisted of 100 trials. The number of hyperparameters for each model and the results of computational cost measurement are provided in Table S4 and The computational costs of each model, including training and sampling wall-clock times and peak GPU memory usage during training and sampling, are summarized in Table S5. Sampling costs were measured for generating 1,000 samples from each trained model.

Table S5, respectively.

**Table S1 Training configurations and hyperparameter search spaces.**

| Setting/Hyperparameter | Synthetic Dataset | NIMH dataset | HCP dataset |
|---|---|---|---|
| **Common Settings** | | | |
| Maximum Epochs | 1000 | 2000 | 1000 |
| Batch Size | 64 | 20 | 64 |
| Early Stopping Patience [epochs] | 30 | | |
| Early Stopping Metric | Val Loss (VAE, DDPM), MMD (WGAN-GP) | | |
| Learning Rate | $[10^{-5}, 10^{-3}]$ | | |
| **Model-Specific Search Space** | | | |
| Latent Dimension (VAE) | $\{2,4,8,16\}$ | $\{4,8,16,32,64\}$ | |
| Latent Dimension (WGAN-GP) | $\{4,8,16,32\}$ | $\{4,8,16,32,64\}$ | |
| Layers (VAE Encoder/Decoder) | Fixed | $[2,5]$ | |
| Layers (WGAN-GP Gen./Critic) | Fixed | $[2,5]$ | |
| U-Net Base Channels (DDPM) | Fixed | $\{8,16,32\}$ | |

VAE=variational autoencoder, WGAN-GP=Wasserstein GAN with gradient penalty, DDPM=denoising diffusion probabilistic models, Gen.=Generator, NIMH = NIMH Healthy Volunteer Dataset, HCP = Human Connectome Project Young Adult S1200 dataset.

**Table S2 Selected hyperparameters for synthetic datasets and models.**

| Datasets | VAE (LR, Latent dim.) | WGAN-GP (Gen. LR, Critic LR, Latent dim.) | DDPM (LR) |
|---|---|---|---|
| BA | $1.0 \times 10^{-3}, 2$ | $2.2 \times 10^{-4}, 1.6 \times 10^{-4}, 32$ | $9.8 \times 10^{-4}$ |
| Planar | $6.6 \times 10^{-4}, 16$ | $6.4 \times 10^{-4}, 1.2 \times 10^{-3}, 16$ | $4.6 \times 10^{-4}$ |
| SBM | $9.9 \times 10^{-4}, 8$ | $1.3 \times 10^{-4}, 3.4 \times 10^{-4}, 32$ | $9.9 \times 10^{-4}$ |
| WS | $8.5 \times 10^{-4}, 2$ | $8.2 \times 10^{-5}, 2.6 \times 10^{-4}, 32$ | $9.4 \times 10^{-4}$ |
| WSF | $1.0 \times 10^{-3}, 2$ | $2.9 \times 10^{-4}, 6.7 \times 10^{-4}, 32$ | $6.3 \times 10^{-4}$ |

VAE=variational autoencoder, WGAN-GP=Wasserstein GAN with gradient penalty, DDPM=denoising diffusion probabilistic models, LR= learning rate, Gen. LR= generator learning rate, Critic LR= critic learning rate, Latent dim.= latent dimension.

**Table S3 Selected hyperparameters for real-world SC dataset and models.**

| | VAE | WGAN-GP | DDPM |
|---|---|---|---|
| **The NIMH Healthy Volunteer Dataset** | | | |
| Learning rate | $2.3 \times 10^{-4}$ | Gen.: $1.3 \times 10^{-4}$, Critic:$2.8 \times 10^{-5}$ | $1.1 \times 10^{-4}$ |
| Latent dim. | 32 | 16 | – |
| Blocks/channels | Encoder: 3, Decoder: 4 | Generator: 2, Critic: 2 | 32 |
| **Human Connectome Project Young-Adult** | | | |
| Learning rate | $1.2 \times 10^{-4}$ | Gen.: $2.4 \times 10^{-5}$, Critic:$1.6 \times 10^{-5}$ | $6.3 \times 10^{-4}$ |
| Latent dim. | 4 | 64 | - |
| Blocks/channels | Encoder: 4, Decoder: 5 | Generator: 2, Critic: 5 | 32 |

VAE=variational autoencoder, WGAN-GP=Wasserstein GAN with gradient penalty, DDPM=denoising diffusion probabilistic models, Latent dim.= latent dimension.

**Table S4 Number of trainable parameters for each model and dataset. Parameter counts are reported in millions (M). The model architectures were identical across runs.**

| Dataset | Model | Params (M) |
|---|---|---|
| Synthetic | VAE | 2.07 |
| | WGAN-GP | 2.09 |
| | DDPM | 2.39 |
| NIMH | VAE | 48.0 |
| | WGAN-GP | 33.9 |
| | DDPM | 9.22 |
| HCP | VAE | 57.7 |
| | WGAN-GP | 4.54 |
| | DDPM | 9.22 |

VAE = variational autoencoder, WGAN-GP = Wasserstein GAN with gradient penalty, DDPM = denoising diffusion probabilistic models, NIMH = NIMH Healthy Volunteer Dataset, HCP = Human Connectome Project Young Adult S1200 dataset.

#### S.6. Computational Cost

The computational costs of each model, including training and sampling wall-clock times and peak GPU memory usage during training and sampling, are summarized in Table S5. Sampling costs were measured for generating 1,000 samples from each trained model.

**Table S5 Computational costs of training and sampling for each model and dataset. Values are**

**reported as mean ± standard deviation (SD) across three runs (runs 2–4). Training and sampling times represent wall-clock time, while the GPU columns indicate peak GPU memory usage.**

| Dataset | Model | Epochs | Training time (s) | Training GPU (MiB) | Sampling time (s) | Sampling GPU (MiB) |
|---|---|---|---|---|---|---|
| BA | VAE | 673 ± 176 | 8.15 ± 4.89 | 67.7 ± 9.3 | 0.086 ± 0.052 | 74.2 ± 9.3 |
| | WGAN-GP | 243 ± 20 | 23.34 ± 0.69 | 70.5 ± 6.9 | 0.058 ± 0.001 | 68.0 ± 7.1 |
| | DDPM | 134 ± 55 | 21.26 ± 8.58 | 1160.9 ± 10.7 | 139.387 ± 0.040 | 2046.1 ± 0.4 |
| Planar | VAE | 173 ± 22 | 1.32 ± 0.53 | 68.2 ± 9.4 | 0.087 ± 0.052 | 74.6 ± 9.4 |
| | WGAN-GP | 174 ± 9 | 11.84 ± 1.44 | 70.4 ± 6.9 | 0.059 ± 0.001 | 67.8 ± 7.1 |
| | DDPM | 173 ± 55 | 12.02 ± 3.54 | 1158.1 ± 11.3 | 139.622 ± 0.084 | 2046.1 ± 0.4 |
| SBM | VAE | 238 ± 78 | 4.06 ± 1.31 | 67.9 ± 9.4 | 0.144 ± 0.003 | 74.4 ± 9.4 |
| | WGAN-GP | 303 ± 227 | 27.56 ± 20.49 | 70.5 ± 6.9 | 0.059 ± 0.001 | 68.0 ± 7.1 |
| | DDPM | 115 ± 55 | 16.59 ± 7.90 | 1160.9 ± 10.7 | 139.509 ± 0.136 | 2046.1 ± 0.4 |
| WS | VAE | 233 ± 150 | 5.73 ± 0.42 | 67.7 ± 9.3 | 0.143 ± 0.002 | 74.2 ± 9.3 |
| | WGAN-GP | 490 ± 32 | 41.24 ± 2.33 | 70.5 ± 6.9 | 0.058 ± 0.001 | 68.0 ± 7.1 |
| | DDPM | 233 ± 150 | 38.11 ± 24.27 | 1160.9 ± 10.7 | 139.451 ± 0.030 | 2046.1 ± 0.4 |
| WSF | VAE | 568 ± 117 | 9.19 ± 2.73 | 67.7 ± 9.3 | 0.087 ± 0.053 | 74.2 ± 9.3 |
| | WGAN-GP | 198 ± 6 | 18.10 ± 1.17 | 70.5 ± 6.9 | 0.058 ± 0.001 | 68.1 ± 7.2 |
| | DDPM | 88 ± 17 | 13.52 ± 3.80 | 1160.9 ± 10.7 | 139.408 ± 0.024 | 2046.1 ± 0.4 |
| NIMH | VAE | 176 ± 8 | 17.84 ± 1.18 | 1179.0 ± 212.2 | 0.108 ± 0.003 | 1046.6 ± 212.2 |
| | WGAN-GP | 383 ± 37 | 91.66 ± 10.22 | 886.1 ± 82.1 | 0.192 ± 0.070 | 564.6 ± 82.3 |
| | DDPM | 115 ± 30 | 24.02 ± 7.59 | 1374.4 ± 43.6 | 834.547 ± 0.369 | 8522.6 ± 0.5 |
| HCP | VAE | 341 ± 29 | 50.81 ± 8.69 | 1416.8 ± 255.3 | 0.106 ± 0.003 | 1244.2 ± 255.7 |
| | WGAN-GP | 512 ± 20 | 108.92 ± 11.37 | 151.6 ± 11.8 | 0.149 ± 0.071 | 139.8 ± 12.2 |
| | DDPM | 154 ± 77 | 350.69 ± 173.72 | 3801.3 ± 40.9 | 836.322 ± 0.136 | 8710.1 ± 41.0 |

VAE = variational autoencoder, WGAN-GP = Wasserstein GAN with gradient penalty, DDPM = denoising diffusion probabilistic models, BA = Barabási-Albert, SBM = stochastic block model, WS = Watts–Strogatz, WSF = weighted scale-free, NIMH = NIMH Healthy Volunteer Dataset, HCP = Human Connectome Project Young Adult S1200 dataset.

#### S.7. Thresholding Results

Table S6 reports the threshold values and edge densities before and after thresholding for each model and run on the synthetic datasets.

**Table S6 Threshold values and edge densities before and after thresholding for synthetic datasets.**

| Dataset | Model | Run | Selected Threshold | Train Dataset Density | Density Before Thresholding | Density After Thresholding |
|---|---|---|---|---|---|---|

| | | | | | | |
|---|---|---|---|---|---|---|
| BA | VAE | run_1 | 0.21 | 0.119 | 1.000 | 0.115 |
| | | run_2 | 0.21 | | 1.000 | 0.117 |
| | | run_3 | 0.21 | | 1.000 | 0.119 |
| | | run_4 | 0.21 | | 1.000 | 0.115 |
| | GAN | run_1 | 0.23 | | 1.000 | 0.120 |
| | | run_2 | 0.26 | | 1.000 | 0.119 |
| | | run_3 | 0.24 | | 1.000 | 0.121 |
| | | run_4 | 0.27 | | 1.000 | 0.118 |
| | DDPM | run_1 | 0.08 | | 0.610 | 0.119 |
| | | run_2 | 0.06 | | 0.592 | 0.119 |
| | | run_3 | 0.27 | | 0.673 | 0.118 |
| | | run_4 | 0.07 | | 0.391 | 0.119 |
| Planar | VAE | run_1 | 0.14 | 0.088 | 1.000 | 0.086 |
| | | run_2 | 0.14 | | 1.000 | 0.088 |
| | | run_3 | 0.14 | | 1.000 | 0.086 |
| | | run_4 | 0.14 | | 1.000 | 0.086 |
| | GAN | run_1 | 0.35 | | 1.000 | 0.089 |
| | | run_2 | 0.34 | | 1.000 | 0.089 |
| | | run_3 | 0.36 | | 1.000 | 0.089 |
| | | run_4 | 0.35 | | 1.000 | 0.089 |
| | DDPM | run_1 | 0.24 | | 0.739 | 0.088 |
| | | run_2 | 0.79 | | 0.719 | 0.089 |
| | | run_3 | 0.31 | | 0.641 | 0.088 |
| | | run_4 | 0.24 | | 0.559 | 0.089 |
| SBM | VAE | run_1 | 0.32 | 0.174 | 1.000 | 0.180 |
| | | run_2 | 0.33 | 0.175 | 1.000 | 0.171 |
| | | run_3 | 0.33 | 0.175 | 1.000 | 0.174 |
| | | run_4 | 0.33 | 0.174 | 1.000 | 0.167 |
| | GAN | run_1 | 0.26 | 0.174 | 1.000 | 0.172 |
| | | run_2 | 0.41 | 0.175 | 1.000 | 0.183 |
| | | run_3 | 0.25 | 0.175 | 1.000 | 0.177 |
| | | run_4 | 0.27 | 0.174 | 1.000 | 0.174 |
| | DDPM | run_1 | 0.51 | 0.174 | 0.365 | 0.174 |
| | | run_2 | 0.89 | 0.175 | 0.525 | 0.173 |
| | | run_3 | 0.29 | 0.175 | 0.701 | 0.176 |

| | | | | | | |
|---|---|---|---|---|---|---|
| | | run_4 | 0.13 | 0.174 | 0.401 | 0.174 |
| WS | VAE | run_1 | 0.24 | 0.095 | 1.000 | 0.095 |
| | | run_2 | 0.27 | | 1.000 | 0.095 |
| | | run_3 | 0.26 | | 1.000 | 0.096 |
| | | run_4 | 0.28 | | 1.000 | 0.094 |
| | GAN | run_1 | 0.29 | | 1.000 | 0.095 |
| | | run_2 | 0.30 | | 1.000 | 0.096 |
| | | run_3 | 0.29 | | 1.000 | 0.095 |
| | | run_4 | 0.29 | | 1.000 | 0.096 |
| | DDPM | run_1 | 0.12 | | 0.577 | 0.095 |
| | | run_2 | 0.22 | | 0.460 | 0.095 |
| | | run_3 | 0.47 | | 0.619 | 0.095 |
| | | run_4 | 0.05 | | 0.452 | 0.092 |
| WSF | VAE | run_1 | 0.13 | 0.223 | 1.000 | 0.221 |
| | | run_2 | 0.13 | | 1.000 | 0.220 |
| | | run_3 | 0.13 | | 1.000 | 0.217 |
| | | run_4 | 0.13 | | 1.000 | 0.214 |
| | GAN | run_1 | 0.11 | | 1.000 | 0.225 |
| | | run_2 | 0.11 | | 1.000 | 0.230 |
| | | run_3 | 0.11 | | 1.000 | 0.230 |
| | | run_4 | 0.11 | | 1.000 | 0.220 |
| | DDPM | run_1 | 0.16 | | 0.669 | 0.224 |
| | | run_2 | 0.13 | | 0.707 | 0.224 |
| | | run_3 | 0.16 | | 0.704 | 0.221 |
| | | run_4 | 0.14 | | 0.615 | 0.222 |

VAE = variational autoencoder, WGAN-GP = Wasserstein GAN with gradient penalty, DDPM = denoising diffusion probabilistic models, BA = Barabási-Albert, SBM = stochastic block model, WS = Watts–Strogatz, WSF = weighted scale-free, NIMH = NIMH Healthy Volunteer Dataset, HCP = Human Connectome Project Young Adult S1200 dataset.

### S.8. Supplementary Results

#### S.8.1. Synthetic Datasets

Individual samples generated by each model across four training runs for the synthetic datasets are shown in Figure S1. Table S7 summarizes MMD-based metrics for each dataset and planarity rate for the Planar dataset across four runs. Table S8 presents the Wasserstein distances between the generated and test distributions of Modularity for the SBM dataset and Small-worldness for the WS dataset.

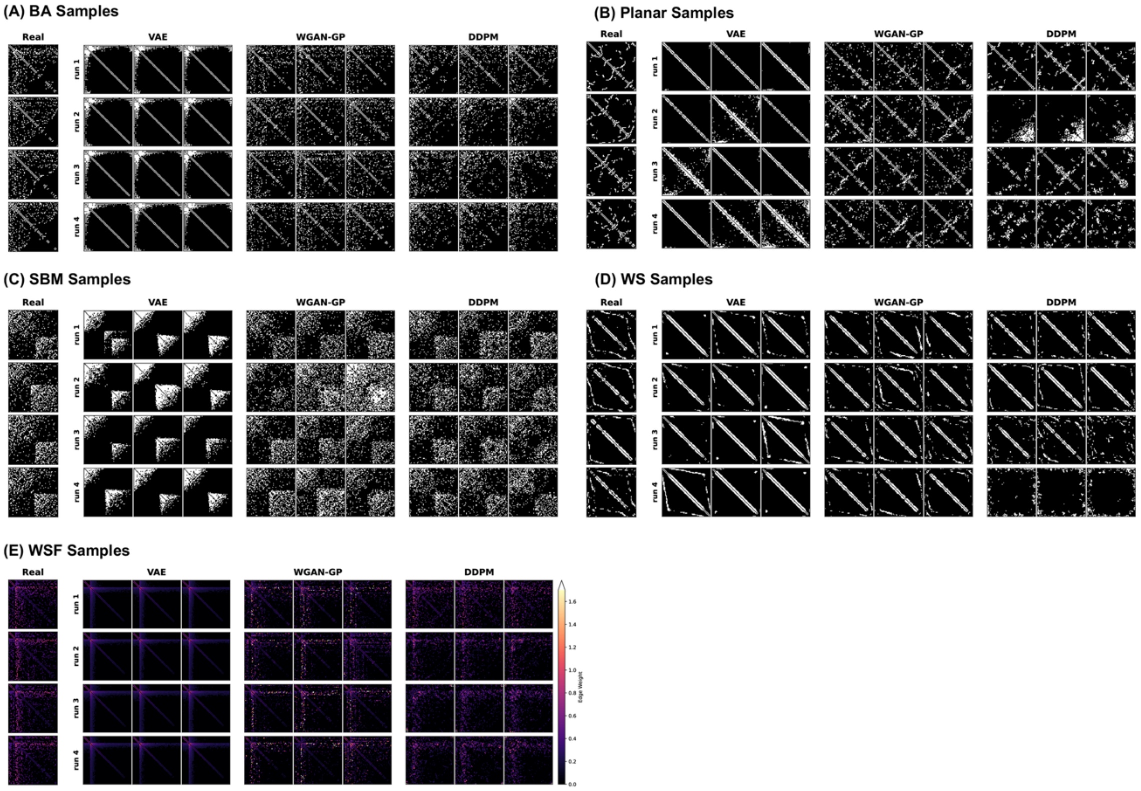


**Figure S1 Individual adjacency matrix analysis across five synthetic datasets (A-E). The real data column represents four randomly selected samples from the test set, while each generated data column represents three randomly selected samples from each of the four training runs. The color bar indicates edge weight, with lighter colors corresponding to higher values. Abbreviations: VAE = variational autoencoder, WGAN-GP = Wasserstein GAN with gradient penalty, DDPM = denoising diffusion probabilistic models, BA = Barabási-Albert, SBM = stochastic block model, WS = Watts–Strogatz, WSF = weighted scale-free.**

144 **Table S7 MMD results for individual training runs on five synthetic datasets.**

**(A) BA**

| | | run 1 | run 2 | run 3 | run 4 |
|---|---|---|---|---|---|
| VAE | Deg. | 1.0 | 1.2 | 1.2 | 1.1 |
| | Clus. | 1.0 | 1.2 | 1.3 | 1.0 |
| | Orbit | 1.7 | 1.9 | 1.9 | 1.7 |
| | Spec. | 0.020 | 0.047 | 0.040 | 0.019 |
| | Emb. | 1.5 | 1.9 | 1.9 | 1.5 |
| GAN | Deg. | 0.68 | 0.68 | 0.69 | 0.66 |
| | Clus. | 0.100 | 0.093 | 0.096 | 0.094 |
| | Orbit | 0.19 | 0.19 | 0.18 | 0.19 |
| | Spec. | 0.0010 | 0.00092 | 0.00069 | 0.0012 |
| | Emb. | 0.33 | 0.34 | 0.33 | 0.33 |
| DDPM | Deg. | 0.79 | 0.77 | 0.76 | 0.80 |
| | Clus. | 0.17 | 0.29 | 0.14 | 0.27 |
| | Orbit | 0.16 | 0.75 | 0.24 | 0.59 |
| | Spec. | 0.0018 | 0.012 | 0.0076 | 0.012 |
| | Emb. | 0.37 | 0.69 | 0.42 | 0.67 |

**(B) Planar**

| | | run 1 | run 2 | run 3 | run 4 |
|---|---|---|---|---|---|
| VAE | Deg. | 0.58 | 0.58 | 0.64 | 0.58 |
| | Clus. | 0.76 | 0.74 | 0.76 | 0.75 |
| | Orbit | 0.061 | 0.033 | 0.047 | 0.034 |
| | Spec. | 0.066 | 0.061 | 0.066 | 0.064 |
| | Emb. | 0.84 | 0.81 | 0.85 | 0.78 |
| | Plan. | 0.16 | 0.13 | 0.17 | 0.13 |
| GAN | Deg. | 0.43 | 0.43 | 0.42 | 0.42 |
| | Clus. | 1.2 | 1.2 | 1.2 | 1.2 |
| | Orbit | 0.90 | 0.92 | 0.91 | 0.95 |
| | Spec. | 0.061 | 0.061 | 0.061 | 0.061 |
| | Emb. | 0.60 | 0.59 | 0.59 | 0.55 |
| | Plan. | 0 | 0 | 0 | 0 |
| DDPM | Deg. | 0.69 | 1.3 | 0.82 | 0.97 |
| | Clus. | 0.97 | 1.5 | 1.1 | 1.3 |
| | Orbit | 0.76 | 1.2 | 0.80 | 0.97 |
| | Spec. | 0.047 | 0.81 | 0.069 | 0.15 |
| | Emb. | 0.64 | 1.3 | 0.80 | 0.91 |
| | Plan. | 0 | 0 | 0 | 0 |

**(C) SBM**

| | | run 1 | run 2 | run 3 | run 4 |
|---|---|---|---|---|---|
| VAE | Deg. | 0.76 | 0.72 | 0.70 | 0.73 |
| | Clus. | 0.99 | 0.88 | 0.84 | 0.93 |
| | Orbit | 1.6 | 1.4 | 1.3 | 1.4 |
| | Spec. | 0.30 | 0.29 | 0.30 | 0.30 |
| | Emb. | 1.1 | 0.91 | 0.88 | 0.92 |
| GAN | Deg. | 0.046 | 0.26 | 0.046 | 0.044 |
| | Clus. | 0.0090 | 0.11 | 0.013 | 0.010 |
| | Orbit | 0.14 | 0.43 | 0.14 | 0.13 |
| | Spec. | 0.0020 | 0.0083 | 0.0019 | 0.0021 |
| | Emb. | 0.11 | 0.45 | 0.11 | 0.10 |
| DDPM | Deg. | 0.073 | 0.039 | 0.066 | 0.065 |
| | Clus. | 0.013 | 0.041 | 0.011 | 0.0096 |
| | Orbit | 0.070 | 0.13 | 0.12 | 0.14 |
| | Spec. | 0.0018 | 0.0033 | 0.0020 | 0.0020 |
| | Emb. | 0.037 | 0.033 | 0.094 | 0.066 |

**(D) WS**

| | | run 1 | run 2 | run 3 | run 4 |
|---|---|---|---|---|---|
| VAE | Deg. | 0.82 | 0.71 | 0.74 | 0.67 |
| | Clus. | 0.59 | 0.55 | 0.52 | 0.54 |
| | Orbit | 0.49 | 0.60 | 0.51 | 0.58 |
| | Spec. | 0.042 | 0.067 | 0.058 | 0.072 |
| | Emb. | 1.0 | 0.98 | 0.98 | 0.98 |
| GAN | Deg. | 1.3 | 1.2 | 1.3 | 1.3 |
| | Clus. | 0.78 | 0.77 | 0.79 | 0.79 |
| | Orbit | 0.35 | 0.35 | 0.35 | 0.37 |
| | Spec. | 0.018 | 0.016 | 0.017 | 0.017 |
| | Emb. | 0.90 | 0.88 | 0.90 | 0.89 |
| DDPM | Deg. | 1.4 | 1.3 | 1.4 | 1.5 |
| | Clus. | 0.83 | 0.67 | 0.96 | 1.3 |
| | Orbit | 0.58 | 0.39 | 1.1 | 1.2 |
| | Spec. | 0.017 | 0.015 | 0.041 | 0.80 |
| | Emb. | 0.97 | 0.96 | 1.00 | 1.1 |

**(E) WSF**

| | | run 1 | run 2 | run 3 | run 4 |
|---|---|---|---|---|---|
| VAE | Deg. | 0.69 | 0.82 | 0.75 | 0.77 |
| | Clus. | 1.0 | 1.1 | 0.92 | 0.95 |
| | Orbit | 1.9 | 2.0 | 1.8 | 1.8 |
| | Spec. | 0.13 | 0.17 | 0.092 | 0.087 |
| | Emb. | 1.7 | 1.9 | 1.5 | 1.5 |
| | Str. | 1.2 | 1.2 | 1.2 | 1.2 |
| GAN | Deg. | 0.32 | 0.31 | 0.32 | 0.34 |
| | Clus. | 0.25 | 0.31 | 0.26 | 0.22 |
| | Orbit | 0.91 | 0.95 | 0.90 | 0.88 |
| | Spec. | 0.012 | 0.010 | 0.010 | 0.014 |
| | Emb. | 0.69 | 0.72 | 0.68 | 0.66 |
| | Str. | 0.13 | 0.16 | 0.13 | 0.13 |
| DDPM | Deg. | 0.37 | 0.35 | 0.38 | 0.36 |
| | Clus. | 0.026 | 0.10 | 0.18 | 0.15 |
| | Orbit | 0.33 | 0.58 | 0.78 | 0.66 |
| | Spec. | 0.0059 | 0.0041 | 0.0021 | 0.0041 |
| | Emb. | 0.31 | 0.55 | 0.59 | 0.53 |
| | Str. | 0.060 | 0.074 | 0.18 | 0.24 |

*Abbreviations*: VAE = variational autoencoder, WGAN-GP = Wasserstein GAN with gradient penalty, DDPM = denoising diffusion probabilistic models, BA = Barabási-Albert, SBM = stochastic block model, WS = Watts–Strogatz, WSF = weighted scale-free, deg. = degree MMD, Clus. = clustering coefficient MMD, Spec. = spectral MMD, Emb. = embedding MMD, Plan. = planarity rate, str. = edge strength MMD.

**Table S8 Wasserstein distances between the generated and test distributions of modularity and small-worldness for the SBM and WS datasets, respectively, across individual training runs.**

**(A) Modularity**

| | run 1 | run 2 | run 3 | run 4 |
|---|---|---|---|---|
| VAE | 0.14 | 0.14 | 0.13 | 0.14 |
| GAN | 0.014 | 0.086 | 0.017 | 0.011 |
| DDPM | 0.017 | 0.070 | 0.034 | 0.044 |

**(B) Small-worldness**

| | run 1 | run 2 | run 3 | run 4 |
|---|---|---|---|---|
| VAE | 1.4 | 1.4 | 1.4 | 1.4 |
| GAN | 0.79 | 0.88 | 0.80 | 0.87 |
| DDPM | 1.0 | 0.80 | 2.2 | 4.3 |

*Abbreviations*: VAE = variational autoencoder, WGAN-GP = Wasserstein GAN with gradient penalty, DDPM = denoising diffusion probabilistic models

#### S.8.2. Structural Connectivity Datasets

Individual samples generated by each model across four training runs for the NIMH and HCP datasets are shown in Figure S2. Table S9 and Table S10 summarize MMD-based metrics for the

NIMH dataset and HCP dataset, respectively, across four training runs and three edge densities (5%, 15%, and 30%). Table S11 and Table S12 present the Wasserstein distances between the generated and test distributions of modularity and small-worldness, respectively, for the SC datasets across four runs and edge densities. Figure S3 visualizes strong-edge reconstruction across four training runs for the NIMH and HCP datasets, showing the overlap between the strongest edges in the mean adjacency matrices of the generated and test datasets. Table S13 summarizes the strong-edge preservation (Jaccard index) and hub preservation (Spearman correlation coefficient) for each training run in both SC datasets.

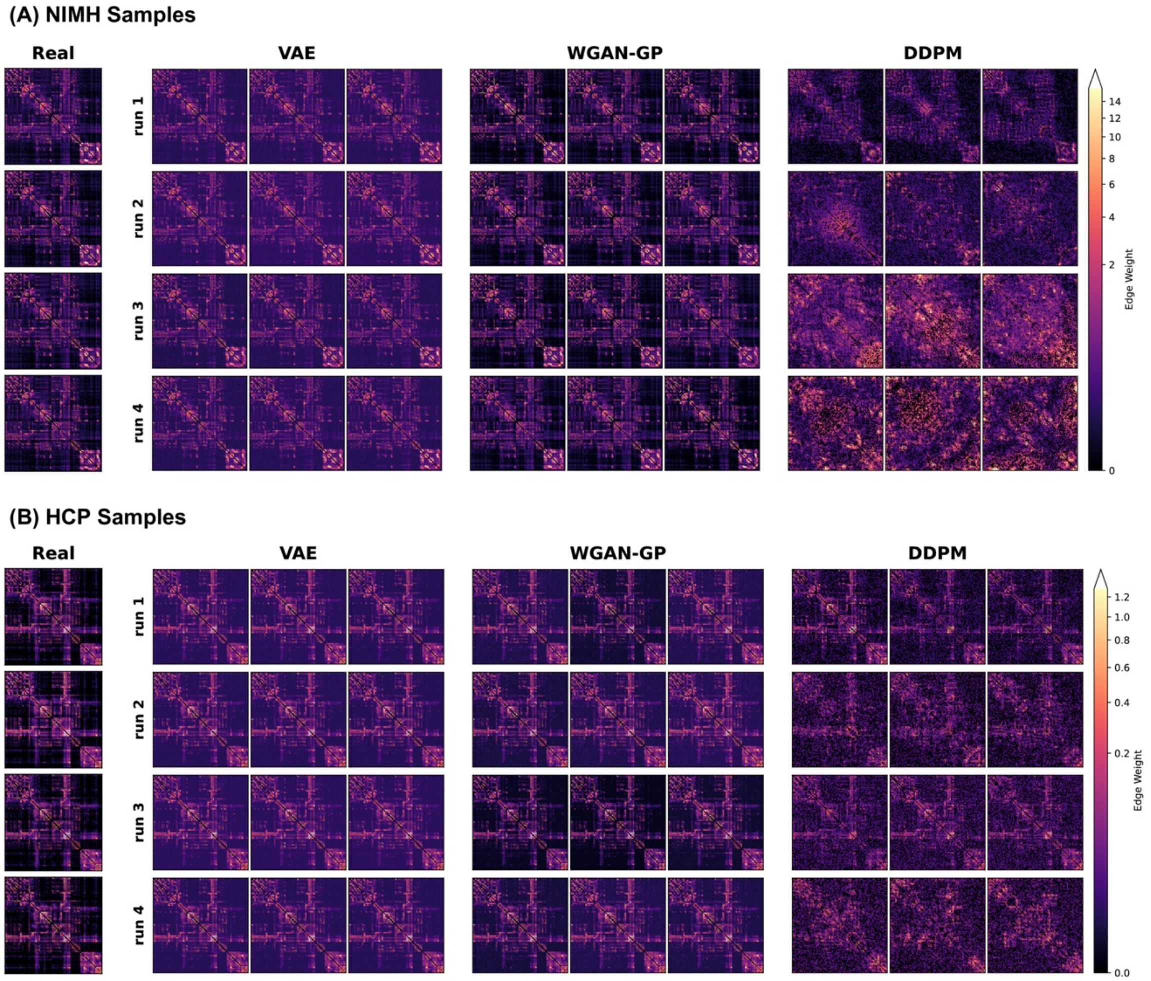


**Figure S2 Individual adjacency matrix analysis across two structural connectivity datasets (A, B). The real data column represents four randomly selected samples from the test set, while each generated data column represents three randomly selected samples from each of the four training runs. The color bar indicates edge weight, with lighter colors corresponding to higher values. *Abbreviations*: VAE = variational autoencoder, WGAN-GP = Wasserstein GAN with gradient penalty, DDPM = denoising diffusion probabilistic models, NIMH = NIMH Healthy Volunteer Dataset, HCP = Human Connectome Project Young Adult S1200 dataset.**

**Table S9 MMD results for individual training runs on the NIMH Healthy Volunteer Dataset.**

| | | d=0.05 | | | | d=0.15 | | | | d=0.3 | | | |
|---|---|---|---|---|---|---|---|---|---|---|---|---|---|
| | | run 1 | run 2 | run 3 | run 4 | run 1 | run 2 | run 3 | run 4 | run 1 | run 2 | run 3 | run 4 |
| VAE | Degree MMD | 0.038 | 0.046 | 0.042 | 0.043 | 0.039 | 0.048 | 0.038 | 0.048 | 0.043 | 0.049 | 0.039 | 0.038 |
| | Clustering MMD | 0.027 | 0.026 | 0.027 | 0.028 | 0.014 | 0.014 | 0.014 | 0.014 | 0.036 | 0.035 | 0.036 | 0.016 |
| | Orbit MMD | 0.38 | 0.39 | 0.32 | 0.31 | 0.20 | 0.28 | 0.20 | 0.22 | 0.39 | 0.43 | 0.41 | 0.42 |
| | Spectral MMD | 0.0039 | 0.0071 | 0.0046 | 0.0045 | 0.013 | 0.018 | 0.011 | 0.015 | 0.018 | 0.023 | 0.016 | 0.021 |
| | Strength MMD | 0.045 | 0.079 | 0.042 | 0.038 | 0.25 | 0.29 | 0.25 | 0.29 | 0.27 | 0.31 | 0.27 | 0.31 |
| | Embedding MMD | 0.17 | 0.14 | 0.12 | 0.15 | 0.13 | 0.21 | 0.15 | 0.20 | 0.40 | 0.48 | 0.38 | 0.45 |
| GAN | Degree MMD | 0.0 | 0.0 | 0.0 | 0.0 | 0.018 | 0.0085 | 0.0073 | 0.011 | 0.011 | 0.0086 | 0.010 | 0.0084 |
| | Clustering MMD | 0.0030 | 0.0 | 0.00038 | 0.0029 | 0.0028 | 0.0039 | 0.0028 | 0.0036 | 0.0051 | 0.0049 | 0.0050 | 0.0047 |
| | Orbit MMD | 0.014 | 0.013 | 0.0062 | 0.015 | 0.0 | 0.0054 | 0.0 | 0.0 | 0.052 | 0.073 | 0.059 | 0.050 |
| | Spectral MMD | 0.0000040 | 0.00027 | 0.0000013 | 0.0010 | 0.00013 | 0.0000055 | 0.000028 | 0.0000028 | 0.00027 | 0.00042 | 0.0012 | 0.00072 |
| | Strength MMD | 0.00080 | 0.0021 | 0.00087 | 0.0019 | 0.017 | 0.018 | 0.016 | 0.019 | 0.018 | 0.020 | 0.018 | 0.022 |
| | Embedding MMD | 0.0065 | 0.0 | 0.0034 | 0.0093 | 0.0 | 0.010 | 0.0040 | 0.0084 | 0.030 | 0.060 | 0.057 | 0.027 |
| DDPM | Degree MMD | 0.29 | 0.41 | 0.43 | 0.49 | 0.077 | 0.068 | 0.055 | 0.084 | 0.097 | 0.039 | 0.029 | 0.020 |
| | Clustering MMD | 0.94 | 1.1 | 1.3 | 1.3 | 0.60 | 0.72 | 0.85 | 0.89 | 0.84 | 0.89 | 0.94 | 1.0 |
| | Orbit MMD | 1.3 | 1.2 | 1.4 | 1.4 | 1.3 | 1.4 | 1.6 | 1.6 | 1.4 | 1.4 | 1.5 | 1.7 |
| | Spectral MMD | 0.040 | 0.064 | 0.041 | 0.087 | 0.058 | 0.066 | 0.068 | 0.081 | 0.057 | 0.083 | 0.13 | 0.14 |
| | Strength MMD | 0.26 | 0.33 | 0.30 | 0.44 | 0.59 | 0.58 | 0.61 | 0.71 | 0.60 | 0.57 | 0.60 | 0.71 |
| | Embedding MMD | 0.79 | 0.86 | 0.77 | 1.0 | 0.19 | 0.15 | 0.086 | 0.32 | 0.39 | 0.71 | 0.83 | 0.80 |

VAE = variational autoencoder, WGAN-GP = Wasserstein GAN with gradient penalty, DDPM = denoising diffusion probabilistic models.

**Table S10 MMD results for individual training runs on the Human Connectome Project Young Adult S1200 dataset.**

| | | d=0.05 | | | | d=0.15 | | | | d=0.3 | | | |
|---|---|---|---|---|---|---|---|---|---|---|---|---|---|
| | | run 1 | run 2 | run 3 | run 4 | run 1 | run 2 | run 3 | run 4 | run 1 | run 2 | run 3 | run 4 |
| VAE | Degree MMD | 0.39 | 0.37 | 0.40 | 0.41 | 0.28 | 0.29 | 0.23 | 0.26 | 0.16 | 0.16 | 0.13 | 0.13 |
| | Clustering MMD | 0.22 | 0.2 | 0.27 | 0.25 | 0.13 | 0.13 | 0.11 | 0.12 | 0.092 | 0.095 | 0.066 | 0.079 |
| | Orbit MMD | 0.51 | 0.59 | 0.53 | 0.52 | 0.48 | 0.49 | 0.48 | 0.46 | 0.55 | 0.55 | 0.47 | 0.52 |
| | Spectral MMD | 0.096 | 0.088 | 0.089 | 0.096 | 0.18 | 0.16 | 0.17 | 0.15 | 0.20 | 0.16 | 0.18 | 0.17 |
| | Strength MMD | 0.73 | 0.76 | 0.75 | 0.73 | 0.75 | 0.77 | 0.77 | 0.74 | 0.76 | 0.79 | 0.78 | 0.75 |
| | Embedding MMD | 0.40 | 0.41 | 0.45 | 0.43 | 0.39 | 0.43 | 0.40 | 0.39 | 0.63 | 0.62 | 0.56 | 0.60 |
| GAN | Degree MMD | 0.0035 | 0.0024 | 0.0027 | 0.0034 | 0.020 | 0.020 | 0.018 | 0.017 | 0.040 | 0.033 | 0.040 | 0.033 |
| | Clustering MMD | 0.0093 | 0.019 | 0.014 | 0.016 | 0.012 | 0.012 | 0.010 | 0.013 | 0.028 | 0.025 | 0.028 | 0.027 |
| | Orbit MMD | 0.15 | 0.14 | 0.14 | 0.17 | 0.20 | 0.21 | 0.18 | 0.23 | 0.39 | 0.37 | 0.38 | 0.37 |
| | Spectral MMD | 0.0019 | 0.0023 | 0.0019 | 0.0022 | 0.0059 | 0.010 | 0.0097 | 0.0097 | 0.0084 | 0.012 | 0.011 | 0.012 |
| | Strength MMD | 0.056 | 0.042 | 0.033 | 0.048 | 0.087 | 0.067 | 0.071 | 0.069 | 0.083 | 0.063 | 0.067 | 0.065 |
| | Embedding MMD | 0.074 | 0.073 | 0.079 | 0.073 | 0.19 | 0.18 | 0.14 | 0.21 | 0.47 | 0.46 | 0.46 | 0.45 |
| DDPM | Degree MMD | 0.22 | 0.52 | 0.28 | 0.30 | 0.015 | 0.036 | 0.016 | 0.070 | 0.10 | 0.13 | 0.11 | 0.19 |
| | Clustering MMD | 0.40 | 1.2 | 0.76 | 1.2 | 0.46 | 0.92 | 0.70 | 0.94 | 0.95 | 1.2 | 1.1 | 1.2 |
| | Orbit MMD | 0.92 | 1.5 | 1.3 | 1.4 | 1.4 | 1.7 | 1.6 | 1.7 | 1.8 | 1.9 | 1.9 | 1.9 |
| | Spectral MMD | 0.017 | 0.13 | 0.024 | 0.059 | 0.041 | 0.12 | 0.052 | 0.11 | 0.038 | 0.18 | 0.10 | 0.16 |
| | Strength MMD | 0.26 | 0.60 | 0.29 | 0.58 | 0.32 | 0.70 | 0.36 | 0.80 | 0.28 | 0.76 | 0.38 | 0.88 |
| | Embedding MMD | 0.65 | 1.0 | 0.80 | 0.30 | 0.27 | 0.92 | 0.43 | 1.3 | 1.8 | 1.9 | 1.8 | 1.9 |

VAE = variational autoencoder, WGAN-GP = Wasserstein GAN with gradient penalty, DDPM = denoising diffusion probabilistic models.

**Table S11 Wasserstein distances between modularity distributions of generated and test data across individual training runs on the NIMH and HCP structural connectivity datasets.**

| | d=0.05 | | | | d=0.15 | | | | d=0.3 | | | |
|---|---|---|---|---|---|---|---|---|---|---|---|---|
| | run 1 | run 2 | run 3 | run 4 | run 1 | run 2 | run 3 | run 4 | run 1 | run 2 | run 3 | run 4 |
| **NIMH** | | | | | | | | | | | | |
| VAE | 0.0048 | 0.0051 | 0.0053 | 0.0048 | 0.015 | 0.015 | 0.013 | 0.014 | 0.023 | 0.020 | 0.021 | 0.019 |
| GAN | 0.0036 | 0.0036 | 0.0047 | 0.0036 | 0.0081 | 0.0084 | 0.0078 | 0.0079 | 0.011 | 0.011 | 0.0086 | 0.0073 |
| DDPM | 0.088 | 0.18 | 0.21 | 0.24 | 0.093 | 0.20 | 0.23 | 0.29 | 0.14 | 0.24 | 0.27 | 0.33 |
| **HCP** | | | | | | | | | | | | |
| VAE | 0.012 | 0.012 | 0.0094 | 0.012 | 0.017 | 0.016 | 0.015 | 0.016 | 0.0085 | 0.0095 | 0.0075 | 0.0090 |
| GAN | 0.0041 | 0.0038 | 0.0042 | 0.0030 | 0.015 | 0.014 | 0.014 | 0.017 | 0.0051 | 0.0042 | 0.0037 | 0.0041 |
| DDPM | 0.054 | 0.12 | 0.090 | 0.11 | 0.053 | 0.14 | 0.079 | 0.12 | 0.072 | 0.18 | 0.12 | 0.18 |

VAE = variational autoencoder, WGAN-GP = Wasserstein GAN with gradient penalty, DDPM = denoising diffusion probabilistic models, NIMH = NIMH Healthy Volunteer Dataset, HCP = Human Connectome Project Young Adult S1200 dataset.

**Table S12 Wasserstein distances between small-worldness distributions of generated and test data across individual training runs on the NIMH and HCP structural connectivity datasets.**

| | d=0.05 | | | | d=0.15 | | | | d=0.3 | | | |
|---|---|---|---|---|---|---|---|---|---|---|---|---|
| | run 1 | run 2 | run 3 | run 4 | run 1 | run 2 | run 3 | run 4 | run 1 | run 2 | run 3 | run 4 |
| **NIMH** | | | | | | | | | | | | |
| VAE | 0.32 | 0.27 | 0.28 | 0.26 | 0.058 | 0.071 | 0.058 | 0.063 | 0.047 | 0.054 | 0.048 | 0.051 |
| GAN | 0.13 | 0.12 | 0.11 | 0.092 | 0.014 | 0.018 | 0.018 | 0.014 | 0.013 | 0.019 | 0.022 | 0.013 |
| DDPM | 2.7 | 2.8 | 2.9 | 3.4 | 0.69 | 0.92 | 0.99 | 1.2 | 0.21 | 0.38 | 0.39 | 0.46 |
| **HCP** | | | | | | | | | | | | |
| VAE | 0.13 | 0.14 | 0.15 | 0.15 | 0.045 | 0.044 | 0.044 | 0.045 | 0.020 | 0.022 | 0.020 | 0.021 |
| GAN | 0.090 | 0.075 | 0.10 | 0.11 | 0.030 | 0.028 | 0.027 | 0.031 | 0.013 | 0.013 | 0.014 | 0.013 |
| DDPM | 1.1 | 2.6 | 1.7 | 2.5 | 0.29 | 0.73 | 0.46 | 0.70 | 0.29 | 0.40 | 0.34 | 0.41 |

VAE = variational autoencoder, WGAN-GP = Wasserstein GAN with gradient penalty, DDPM = denoising diffusion probabilistic models, NIMH = NIMH Healthy Volunteer Dataset, HCP = Human Connectome Project Young Adult S1200 dataset.

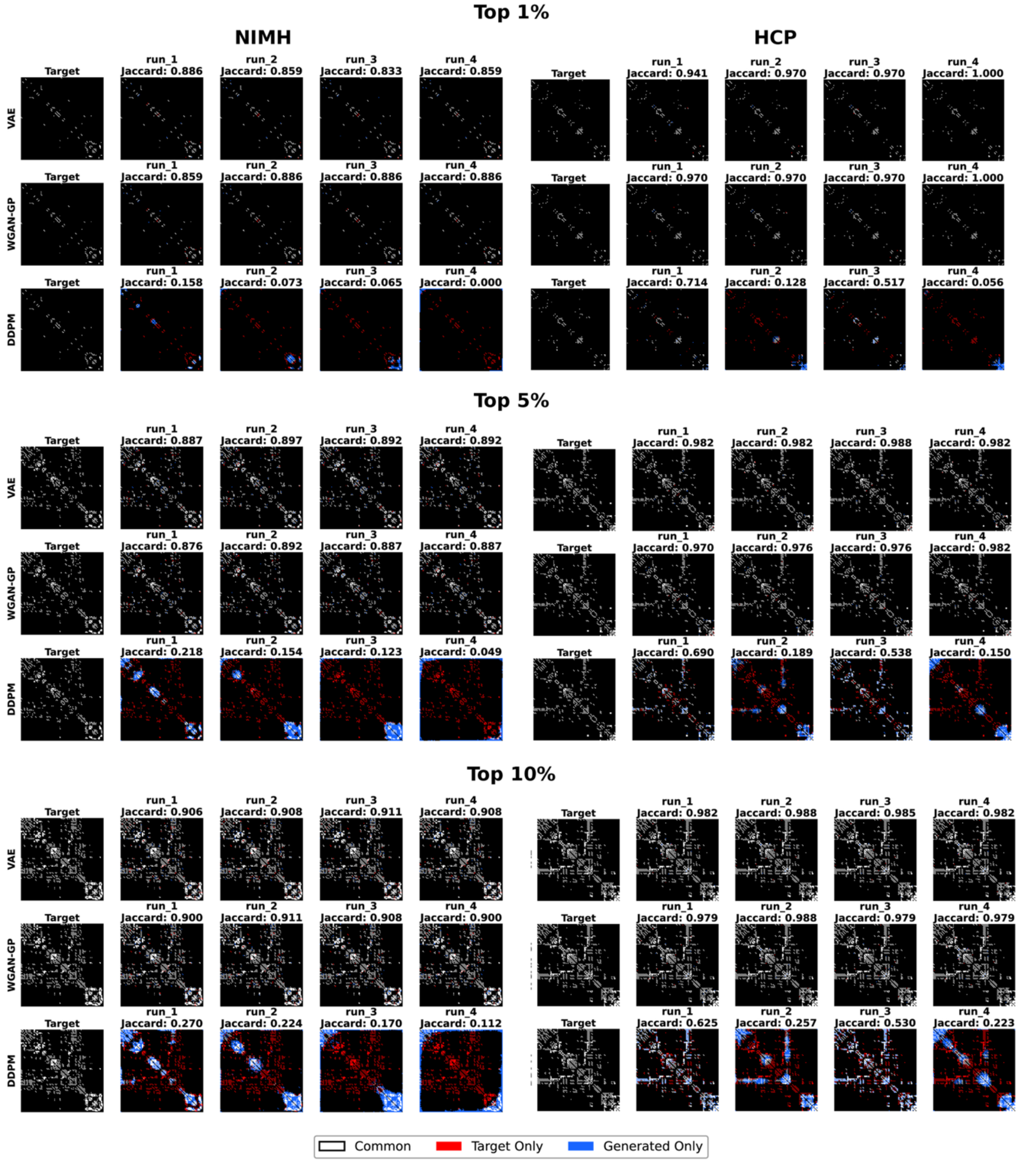


**Figure S3 Strong edge reconstruction between generated graphs and the test datasets for the NIMH and HCP datasets. Results show the top 1%, 5%, and 10% strongest edges in the mean adjacency matrices, arranged from top to bottom. Each row represents a generative model, and each column represents the test dataset (target) or one of the four training runs. White indicates common edges, red indicates target-only edges, and blue indicates generated-only edges. The Jaccard index of strong edge overlap is shown above each generated matrix. Abbreviations: VAE = variational autoencoder, WGAN-GP = Wasserstein GAN with gradient penalty, DDPM = denoising diffusion probabilistic models, NIMH = NIMH Healthy Volunteer**

**Dataset, HCP = Human Connectome Project Young Adult S1200 dataset.**

**Table S13 Strong-edge and hub preservation for each run in the NIMH and HCP datasets.**

| Dataset | model | run | Jaccard index | | | Spearman $\rho$ |
|---|---|---|---|---|---|---|
| | | | Top 1% | Top 5% | Top 10% | |
| NIMH | VAE | 1 | 0.886 | 0.887 | 0.906 | 0.991*** |
| | | 2 | 0.859 | 0.897 | 0.908 | 0.988*** |
| | | 3 | 0.833 | 0.892 | 0.911 | 0.992*** |
| | | 4 | 0.859 | 0.892 | 0.908 | 0.991*** |
| | WGAN-GP | 1 | 0.859 | 0.876 | 0.900 | 0.992*** |
| | | 2 | 0.886 | 0.892 | 0.911 | 0.989*** |
| | | 3 | 0.886 | 0.887 | 0.908 | 0.992*** |
| | | 4 | 0.886 | 0.887 | 0.900 | 0.990*** |
| | DDPM | 1 | 0.158 | 0.218 | 0.270 | 0.297** |
| | | 2 | 0.073 | 0.154 | 0.224 | 0.419*** |
| | | 3 | 0.065 | 0.123 | 0.170 | 0.235* |
| | | 4 | 0.000 | 0.049 | 0.112 | 0.125 |
| HCP | VAE | 1 | 0.941 | 0.982 | 0.982 | 0.999*** |
| | | 2 | 0.970 | 0.982 | 0.988 | 0.999*** |
| | | 3 | 0.970 | 0.988 | 0.985 | 0.999*** |
| | | 4 | 1.000 | 0.982 | 0.982 | 0.999*** |
| | WGAN-GP | 1 | 0.970 | 0.970 | 0.979 | 0.999*** |
| | | 2 | 0.970 | 0.976 | 0.988 | 0.999*** |
| | | 3 | 0.970 | 0.976 | 0.979 | 0.999*** |
| | | 4 | 1.000 | 0.982 | 0.979 | 0.999*** |
| | DDPM | 1 | 0.714 | 0.690 | 0.625 | 0.928*** |
| | | 2 | 0.128 | 0.189 | 0.257 | 0.577*** |
| | | 3 | 0.517 | 0.538 | 0.530 | 0.900*** |
| | | 4 | 0.056 | 0.150 | 0.223 | 0.445*** |

* $p < 0.05$, ** $p < 0.01$, *** $p < 0.001$.

VAE = variational autoencoder, WGAN-GP = Wasserstein GAN with gradient penalty, DDPM = denoising diffusion probabilistic models, NIMH = NIMH Healthy Volunteer Dataset, HCP = Human Connectome Project Young Adult S1200 dataset.
